# IL-7-mediated expansion of autologous lymphocytes increases CD8^+^ VLA-4 expression and accumulation in glioblastoma models

**DOI:** 10.1101/2024.04.01.587634

**Authors:** Kirit Singh, Kelly M. Hotchkiss, Sarah L. Cook, Pamy Noldner, Ying Zhou, Eliese M. Moelker, Chelsea O. Railton, Emily E. Blandford, Bhairavy J. Puviindran, Shannon E. Wallace, Pamela K. Norberg, Gary E. Archer, Beth H. Shaz, Katayoun Ayasoufi, John H. Sampson, Mustafa Khasraw, Peter E. Fecci

## Abstract

The efficacy of T cell-activating therapies against glioma is limited by an immunosuppressive tumor microenvironment and tumor-induced T cell sequestration. We investigated whether peripherally infused non-antigen specific autologous lymphocytes (ALT) could accumulate in intracranial tumors. We observed that non-specific autologous CD8^+^ ALT cells can indeed accumulate in this context, despite endogenous T cell sequestration in bone marrow. Rates of intratumoral accumulation were markedly increased when expanding lymphocytes with IL-7 compared to IL-2. Pre-treatment with IL-7 ALT also enhanced the efficacy of multiple tumor-specific and non-tumor-specific T cell-dependent immunotherapies against orthotopic murine and human xenograft gliomas. Mechanistically, we detected increased VLA-4 on mouse and human CD8^+^ T cells following IL-7 expansion, with increased transcription of genes associated with migratory integrin expression (*CD9)*. We also observed that IL-7 increases *S1PR1* transcription in human CD8^+^ T cells, which we have shown to be protective against tumor-induced T cell sequestration. These observations demonstrate that expansion with IL-7 enhances the capacity of ALT to accumulate within intracranial tumors, and that pre-treatment with IL-7 ALT can boost the efficacy of subsequent T cell-activating therapies against glioma. Our findings will inform the development of future clinical trials where ALT pre-treatment can be combined with T cell-activating therapies.

**Brief Summary:** T cell immunotherapies are limited by few T cells in glioma. Adoptively transferred lymphocytes expanded with IL-7 exhibit increased VLA-4 expression and accumulate in tumors.

## Introduction

T cell-activating immunotherapies, such as immune checkpoint blockade (ICB), require functional T cells at the tumor site to be effective (1–3). Unfortunately, intracranial tumors are harbored within the immunologically distinct central nervous system (CNS) and sit shielded by an immunosuppressive tumor micro-environment (TME) and the blood-brain barrier (BBB) (4). We have shown that CNS tumors, such as glioblastoma, also possess the unique ability to sequester naïve T cells within bone marrow (5), limiting their access to the TME. Likewise, we have more recently described a central role for tumor-associated macrophages (TAM) in driving glioma-infiltrating T cells towards a terminally exhausted state (6), limiting their antitumor capacities. Accordingly, high-grade gliomas possess low numbers of tumor infiltrating lymphocytes (TILs (7)) and have proven minimally responsive to ICB in clinical trials (8–10). This paucity of functional TILs will similarly limit the therapeutic efficacy of newer synthetic therapies that rely on endogenous T cells, including brain bi-specific T cell engagers (BRiTEs) (11). Strategies for enhancing functional T cell presence within CNS tumors are needed.

Currently, strategies to increase T cell numbers in tumor include expanding the T cell pool intratumorally; encouraging T cell trafficking from the periphery; or adding T cells systemically. Each of these face major challenges. Expansion of local lymphocytes is limited by subsequent severe exhaustion amidst tumor-infiltrating CD8^+^ T cells, making them less proliferative to stimulation (12). Recruitment of peripheral lymphocytes can be boosted by immunostimulatory cytokines (e.g., IL-7 (13) or IL-12 (14)), but peripheral cytokine administration can induce profound systemic toxicity (15–17). Local administration of cellular therapies or immunostimulatory cytokines is an alternative but requires an invasive procedure, limiting the potential for repeat administrations (18, 19). Ultimately, peripheral infusion of unmodified or CAR-T cells is an established and easily repeatable method of adding lymphocyte populations to tumors within the CNS (20, 21). Modifying this approach, then, we asked whether peripherally infused non-antigen specific autologous lymphocytes would accumulate in established CNS tumors and synergize with T cell-activating/ICB therapies.

Currently, cellular therapies such as adoptive lymphocyte transfer (ALT) typically use cytokines such as Interleukin-2 (IL-2) or IL-7 to support *ex vivo* T cell expansion (22). IL-2 skews lymphocytes towards a terminally differentiated T effector (T_EFF_) phenotype. While T_EFF_ cells do express migratory integrins, they often have a short half-life once removed from culture (23, 24). IL-7, in turn, promotes homeostatic T cell expansion and expands the central memory T cell (T_CM_) pool, leading to greater T cell persistence in vivo (25, 26). Peripheral administration of long-acting IL-7 has been shown to enhance T cell accumulation in murine glioma and boosts the efficacy of T cell engagers against solid tumors (13, 27). We therefore sought to evaluate which growth factor might produce a cellular product with favorable BBB-penetrance and enhanced anti-tumor efficacy in models of established intracranial glioma.

Herein, we report that peripherally administered T cells (ALT) lacking antigen specificity accumulate in established murine intracranial tumors, despite tumor-imposed T cell sequestration in bone marrow. Rates of intratumoral accumulation are markedly increased when T cells are expanded with IL-7 (IL-7-ALT), which also increases the efficacy of accompanying immunotherapeutic platforms. We find that IL-7 increases expression of the migratory integrin Very Late Antigen-4 (VLA-4) on both mouse and human CD8^+^ T cells, while VLA-4 blockade abrogates the enhanced accumulation of IL-7-ALT in tumors. Transcriptional analysis of hPBMCs expanded with IL-7 reveals upregulation of genes involved in integrin expression and tumor infiltration (*CD9, VLA-6, EPHA4).* Also upregulated is *S1PR1* (sphingosine-1-phosphate receptor 1, also described as S1P1), increased levels of which are known to avert T cell sequestration and license immunotherapeutic responses to glioma (5). These results provide a route to optimizing ALT expansion, as well as to using ALT as an immunotherapeutic adjunct.

## Results

### IL-7-expanded CD8^+^ T cells demonstrate increased accumulation within orthotopic glioblastoma models despite endogenous T cell sequestration in bone marrow

Though peripherally administered, activated autologous T cells have been shown to cross the BBB under physiologic conditions (28), their ability to enter the CNS in the setting of tumor-directed T cell sequestration was unclear. Previous work by our group established that sequestration predominantly impacts naïve T cells (endogenous or ALT), while memory T cells are largely unperturbed (5). We therefore began by examining the rate of CNS tumor accumulation for *ex vivo*-expanded non-specific autologous T cells.

First, we developed T cell expansion processes that skewed phenotypes towards effector (T_EFF,_ CD44^high^CD62L^low^) or central memory (T_CM,_ CD44^high^CD62L^high^) subsets. Expansion of T cells with either 100IU/mL IL-2 or 20ng/mL IL-7 was chosen to enrich for CD8^+^ T_EFF_ or CD8^+^ T_CM_ phenotypes, respectively, based on prior experience and published protocols (22, 29, 30). Following expansion, flow cytometry was performed to assess the makeup of the ALT product pre-administration (gating and expansion strategy in Figure S1, A and B). *Ex vivo* expansions with both IL-2 and IL-7 yielded a cellular product of >95% CD3^+^ cells (Figure S1C) which predominantly consisted of CD8^+^ T cells (∼80%, Figure S1D). The IL-2-ALT product was preferentially skewed towards T_EFF_ (Figure S1E) while the IL-7-ALT product skewed towards T_CM_ (Figure S1F). Few naïve CD8^+^ T cells (T_N_, CD44^low^CD62L^high^) were present at the end of expansion with either cytokine (Figure S1G).

To establish the trafficking capabilities of non-specific CD45.1^+^ T cells in the setting of tumor, we intracranially implanted CT2A stably transfected with the glioma-specific neoantigen Epidermal Growth Factor Receptor variant III (EGFRvIII) into congenic CD45.2^+^ C57/BL6 mice. CT2A is syngeneic on the C57 background and exhibits high heterogeneity, stemness, and resistance to therapy (31). We have also previously described tumor-induced T cell sequestration in bone marrow using this model (5).

After tumors were well-established at 13 days, mice were given a single ALT of either IL-2 or IL-7-expanded CD45.1^+^ T cells (Figure 1A). Following ALT, tumor and bone marrow were collected daily from separate mice over 4 days. Flow cytometry was used to measure changes to the viable exogenous (CD45.1^+^) and endogenous (CD45.2^+^) CD3^+^ T cell compartment (flow gating in Figure S2A). By 48-hours post administration, we detected an increased proportion of the IL-7-ALT dose within tumor compared to IL-2-ALT. (Figure 1B, IL-7-ALT vs IL-2-ALT in tumor, p<0.05*). No significant difference between dose proportions in bone marrow was seen at the same timepoint (Figure 1B, p=0.8172). Examination of intratumoral T cells over time found that accumulation rates for IL-7-ALT CD8^+^ but not CD4^+^ lymphocytes were significantly greater than for IL-2-ALT. In both groups, endogenous CD8^+^ T cell populations in tumor declined (time course shown for CD4^+^ T cells in Figure 1C, CD8^+^ T cells in Figure 1D. 48-hour IL-7-ALT CD8^+^ T cells 31.24x vs IL-2-ALT 4.14x p<0.01**, 72-hour IL-7-ALT CD8^+^ T cells 42.2x vs IL-2-ALT 5.88x, p<0.01**). Counts of IL-7-ALT CD8^+^ T cells in tumor were also increased compared to IL-2-ALT throughout, though differences were non-significant (Figure S2B).

**Figure 1.**
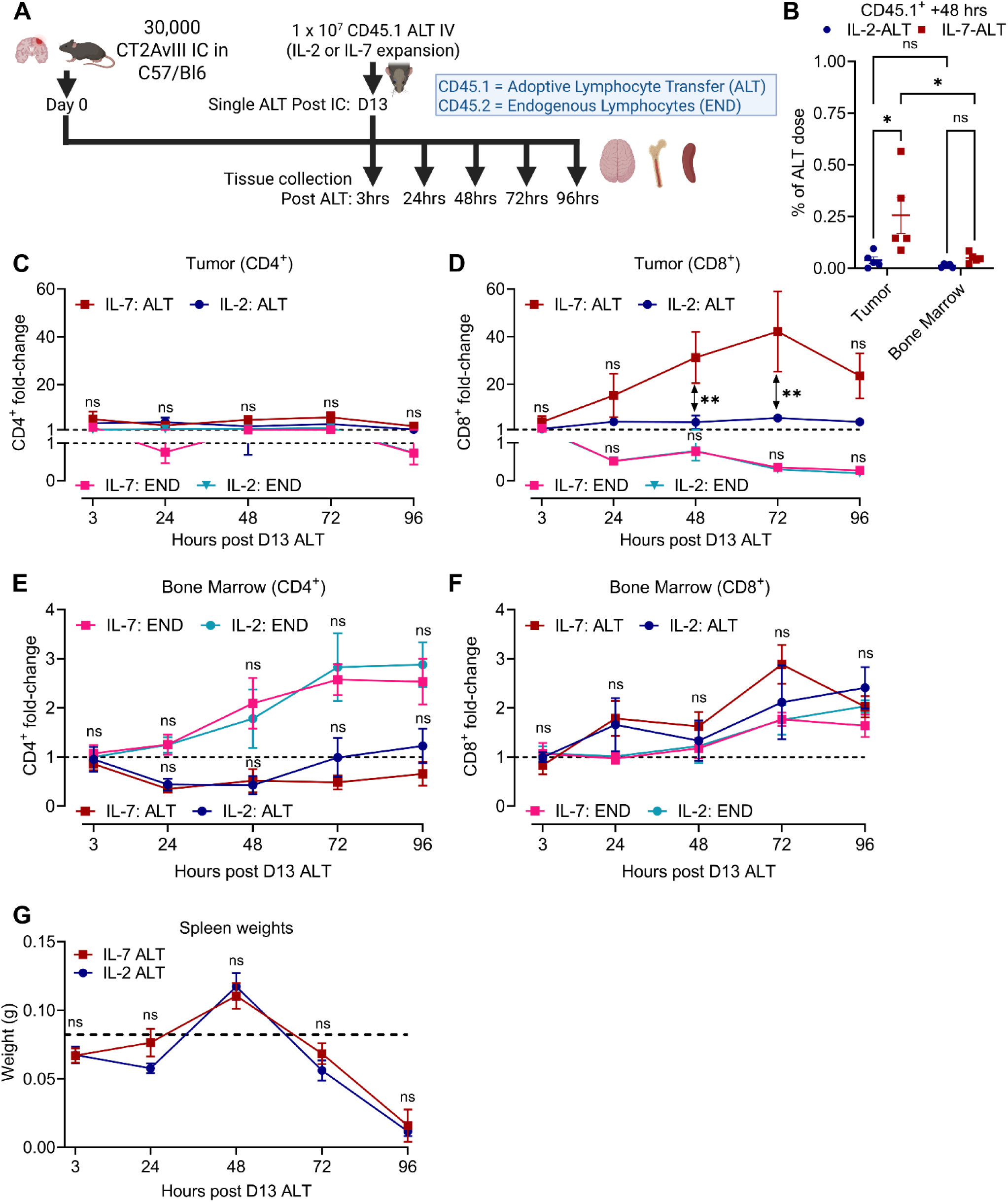
IL-7-ALT CD8^+^ cells demonstrate increased accumulation within orthotopic glioblastoma models despite endogenous T cell sequestration in bone marrow. **(A)** Study overview (n=4-5 mice/group/time point for all graphs). **(B)** Percentage of IL-2-ALT and IL-7-ALT CD45.1^+^ dose detected in tumor by 48-hours post administration (CD4^+^ & CD8^+^ T cells). **(C)** Fold-change relative to Day 0 of exogenous CD45.1^+^CD4^+^ T cell entry and **(D)** CD45.1^+^CD8^+^ T cell entry into brain tumors. **(E)** Fold-change relative to Day 0 of exogenous CD45.1^+^CD4^+^ T cell entry and **(F)** CD45.1^+^CD8^+^ T cell entry into bone marrow. **(G)** Time course of spleen sizes following ALT. Weights from control mice represented by dashed line (0.0823g, n=5). Dashed line in C-F represents baseline (i.e., 1x). Statistical analyses performed via two-way ANOVA and data presented as mean ± SEM unless otherwise specified. Experimental outline generated using BioRender.com.

Simultaneously, increases in endogenous CD4^+^ and CD8^+^ T cell populations in bone marrow were seen across groups over time (CD4^+^ T cells in Figure 1E, CD8^+^ T cells in Figure 1F). Greater fold-change increases in endogenous CD4^+^ T cell accumulation in bone marrow were observed compared to CD8^+^ T cells, in keeping with previous descriptions of tumor-induced CD4^+^ T cell sequestration (endogenous CD4^+^ fold-change of 2.69x vs CD8^+^ T cells fold change of 1.76x at 72-hr observation point (5, 32, 33)). ALT entry into bone marrow was also observed, although this was restricted to the CD8^+^ population. Rates of ALT accumulation in bone marrow did not differ between expansion conditions (Figure 1F, non-adjusted IL-7-ALT CD8^+^ T cell counts in Fig. S2C). Increases in spleen weights were also seen following ALT, before a subsequent precipitous decline below baseline (time course in Figure 1G). We concluded then that non-antigen specific, exogenously administered CD8^+^ T cells can indeed accumulate within glioma despite tumor-imposed T cell sequestration, with IL-7-ALT accumulating intratumorally at a greater rate than IL-2-ALT.

### IL-7-ALT synergizes with T cell-centric immunotherapies in orthotopic glioma models

We next examined whether IL-2 or IL-7 ALT would each enhance survival when combined with T cell-activating immunotherapies. In a first set of experiments, we combined each ALT with EGFRvIII-BRiTE in a U87 xenograft model stably transfected with EGFRvIII (U87vIII). EGFRvIII-BRiTE is a bi-specific T cell engager possessing specificity for both the CD3 receptor and the glioma-specific antigen EGFRvIII (34). As EGFRvIII-BRiTE is fully humanized, we evaluated efficacy in a NOD scid gamma (NSG) mouse model reconstituted with either IL-7- or IL-2-expanded human (h)PBMCs. We observed that IL-7 ALT yielded a significant but modest survival benefit when combined with BRiTE, compared to the IL-2-ALT-BRITE combination (Figure 2A, p<0.05*). Interestingly, no differences between the two modalities were observed in vitro (Figure 2B).

**Figure 2.**
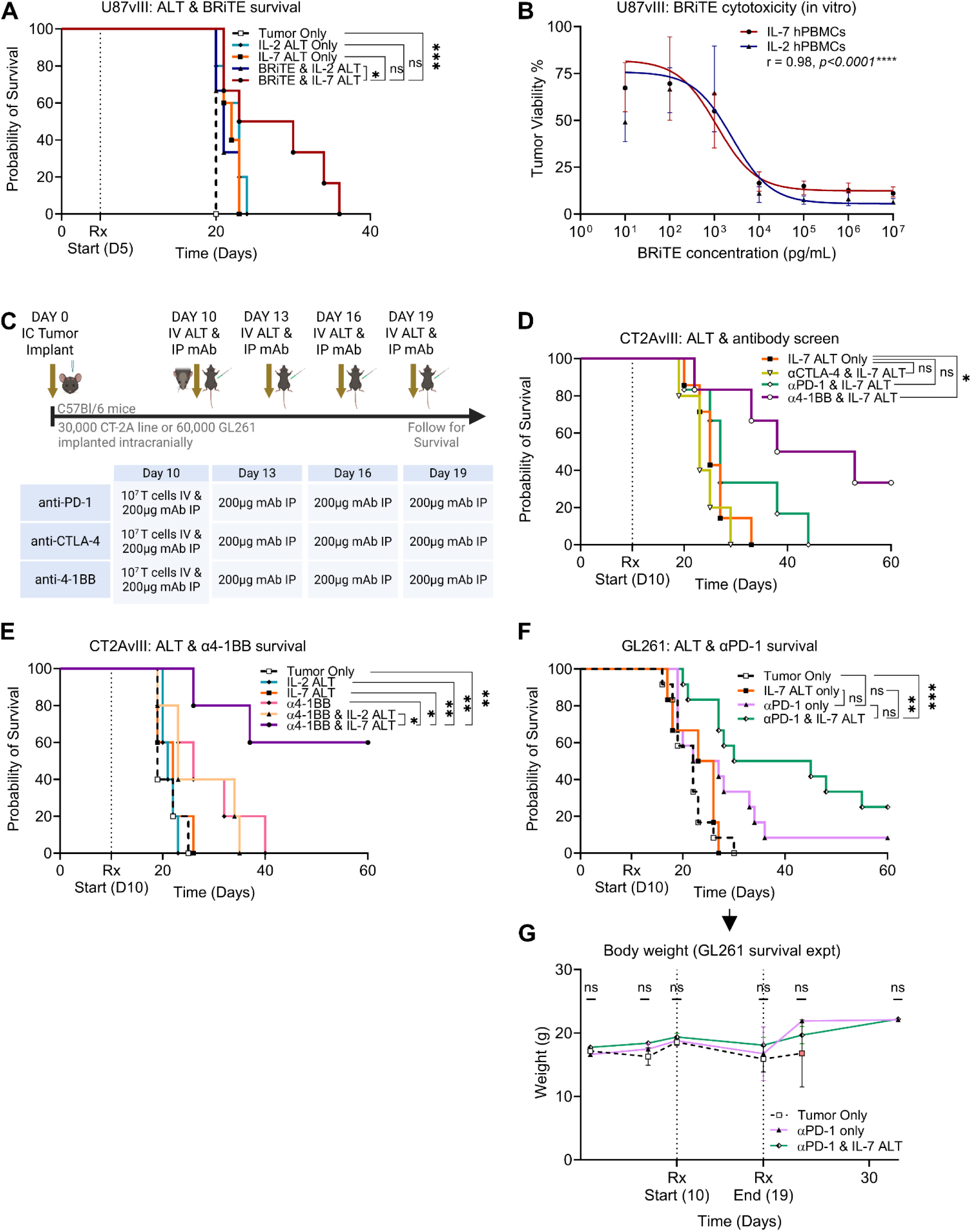
IL-7-ALT synergizes with specific and non-specific T cell-activating/checkpoint blockade therapies in orthotopic glioma models. **(A)** Evaluation of IL-7 and IL-2 ALT combined with hCD3:EGFRvIII BRiTE in NSG mice. Mice (n=5-6 /group) were implanted with U87vIII and treated with 5 x 10^6^ IV hPBMCs on day 5, with serial IV BRiTE (50µg) on days 5-9. **(B)** Cytotoxicity assay using BRiTE co-cultured with tumor cells (U87vIII) and hPBMCs expanded with IL-2 or IL-7. Non-linear fitted dose-response curves shown with SEM (n=10-12/group, Pearson Correlation Coefficient r = 0.98, p<0.0001****). **(C & D)** Study overview and findings of a screening approach to identify the best combinatorial mAb approach with ALT (n=5-7/group). **(E)** Survival of combination α4-1BB & IL-7 ALT therapy compared to monotherapy controls treated using same regimen in **(C)** (n=5/group). **(F)** C57BL/6 mice implanted with 6 x 10^4^ GL261 and treated using same regimen in **(C)** with αPD-1 (n=6-12/group, pooled data across two experiments). **(G)** Monitoring of bodyweight throughout the treatment period and for two weeks after for survival experiment in **(F)** (n=6-7/group). Comparison via multiple unpaired t-tests and data presented as mean ± SEM. Survival comparisons performed via a log-rank (Mantel-Cox) chi-squared test. Experimental outlines generated using BioRender.com.

Having observed a survival benefit with the combination of IL-7-ALT + BRITE, we looked to see how successfully IL-7-ALT might also combine with various checkpoint or T cell activity modifying therapies that typically have limited or no success in CT2A models (outline in Figure 2C). After CT2AvIII tumors established for 10 days, animals were treated with IL-7 ALT alone, or in combination with αCTLA-4 (antagonist), αPD-1 (antagonist) or α4-1BB (agonist) monoclonal antibodies. We observed a small trend toward prolonged survival when combining IL-7 ALT with αPD-1 therapy (2/6 mice surviving > 30 days, 0/6 with long term overall survival), but a significant increase in survival when combining IL-7-ALT + α4-1BB compared to IL-7-ALT only (Figure 2D, p<0.05*). We therefore selected the IL-7-ALT + α4-1BB combination approach for further experiments in this model and performed controlled survival studies using the same timing schematic as above (Figure 2E). We again found that IL-7-ALT + α4-1BB therapy yielded a significant survival benefit, this time compared to combination IL-2-ALT and α4-1BB (p<0.05*); α4-1BB alone (p<0.05*); either IL-2 or IL-7-ALT only (both p<0.01**); or to untreated controls (p<0.01**). Interestingly, while combination IL-7 ALT + αPD-1 therapy exhibited a limited survival benefit in the CT2AvIII glioma model (Figure 2D), greater efficacy was observed in mice bearing more immunogenic GL261 gliomas (35) (Figure 2F).

We likewise evaluated whether adding IL-7 ALT to clinically relevant checkpoint blockade therapy would result in toxicity. Body weights during and for two weeks following treatment were recorded in the GL261-bearing mice whose survival is depicted in Figure 2F. No differences between groups were seen (Figure 2G). To evaluate toxicity in greater detail, we performed acute toxicity studies in healthy mice receiving IL-7 + αPD-1 therapy, comparing against sham controls (outline shown in Figure S3A). Body weights again remained stable throughout (Figure S3B). Five days after treatment-end, blood, liver tissue, spinal cord and brain were taken to assess for liver function, changes in cellular architecture and potential demyelination. No significant difference in serum chemistry was seen between groups (Figure S3C). Review of brain and spinal cord by a board-certified veterinary pathologist also found no differences in terms of histological features, cellular architecture, or demyelination (representative photomicrographs in Figure S3, D and E). Regarding liver, no histopathologic abnormalities related to lymphocytic infiltration were noted in the sham group, though presumed peri-mortem acute hepatocellular apoptosis/necrosis without associated inflammation was seen in 2 of 5 animals. In the combination group, pathologist review identified small numbers of randomly distributed foci (1 to 4 per animal) of acute to subacute hepatocellular apoptosis/necrosis associated with a mixed lymphocytic infiltrate (representative photomicrographs in Figure S3F). While these findings were considered to be treatment-related effects, they were considered unlikely to indicate systemic toxicity given the unchanged liver function tests.

### IL-7-expanded CD8^+^ T cells accumulating in tumor exhibit both central and effector memory phenotypes

Frequent antigenic stimulation upregulates trafficking molecules on T_CM_ cells (36, 37), while T_EFF_ cells also upregulate migratory ligands such as LFA-1 and VLA-4, enhancing endothelial binding following activation (23, 24). We therefore evaluated the phenotypic makeup of exogenous T cells that entered the brain following ALT, as well as their expression of migratory integrins.

To begin, we intravenously administered CD45.1^+^ IL-2-ALT or IL-7-ALT to CD45.2^+^ mice with established (15-day) CT2AvIII tumors. Tumors were collected 3- and 48-hours post ALT (outline in Figure 3A). Again, we observed significantly enhanced accumulation of IL-7-ALT CD8^+^ T cells compared to IL-2-ALT (Figure 3B, 3-hrs p<0.01**, 48-hrs p<0.001***). Similar to previous findings, ALT in tumor primarily consisted of CD8^+^ T cells (Figure 3C, CD8^+^ vs CD4^+^ T cells, p<0.0001****). By the 48-hour timepoint, IL-7-ALT T cells made up approximately ∼20% of all T cells in tumor, (Figure 3D, mean 18.8%, SD ±4.3%, n=5), while the fraction of T cells that consisted of IL-2-ALT in tumor was ∼1% (mean 1.7%, SD ±1.5%, n=4).

**Figure 3.**
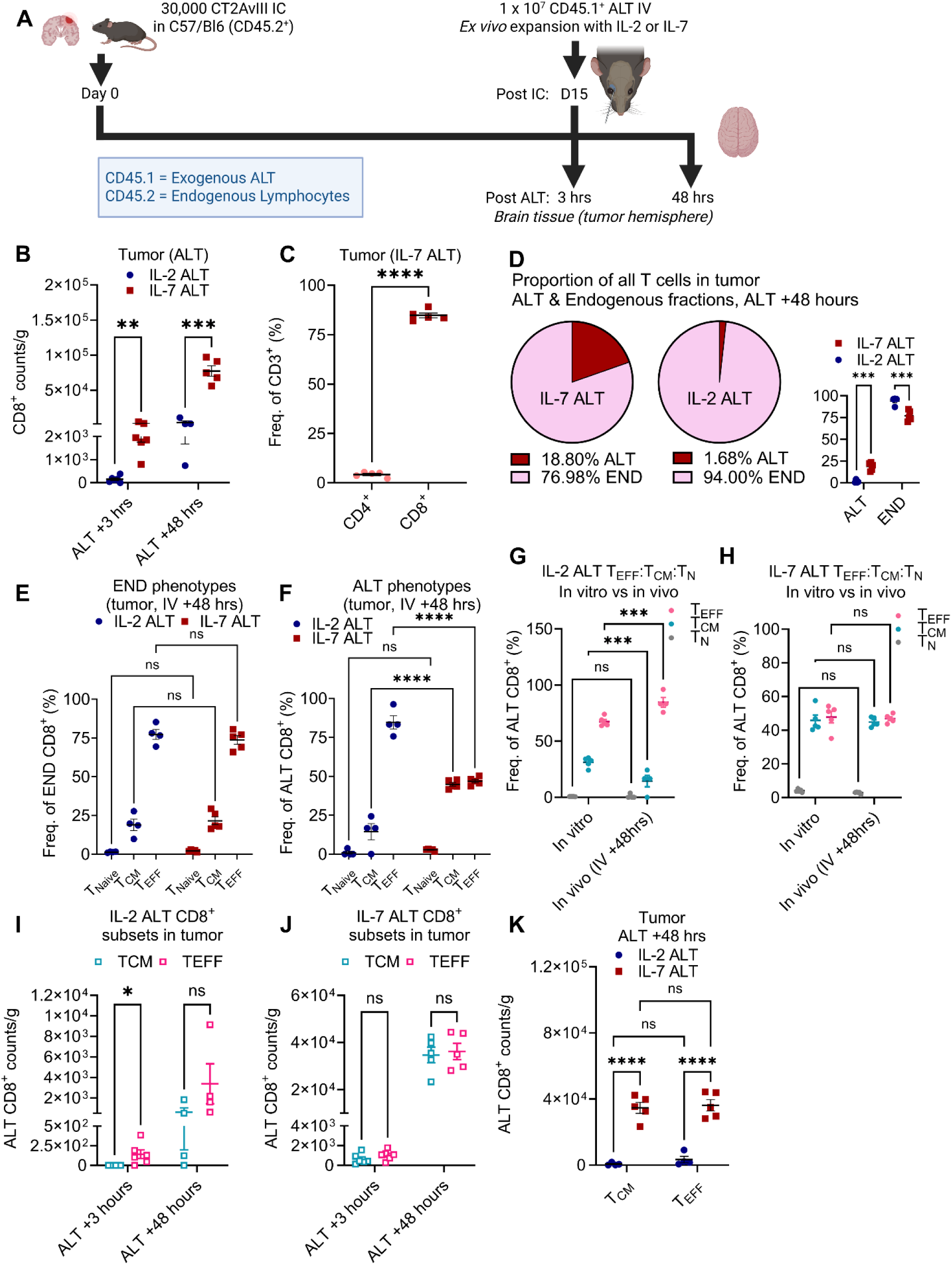
IL-7-expanded CD8^+^ T cells accumulating in tumor consist of both central and effector memory phenotypes. **(A)** Study overview in C57BL/6 mice (n=4-6/group for all graphs). **(B)** Weight-adjusted counts of CD45.1^+^CD8^+^ T cells 3- and 48-hours following ALT in tumor (multiple unpaired t-tests shown). **(C)** Proportion of IL-7-ALT cells in tumor that are CD4^+^ or CD8^+^ T cells (unpaired t-test). **(D)** Proportions of exogenous ALT to endogenous T cells in tumor 48-hours following administration, with paired dot-plot showing individual values across groups. **(E)** Comparison of Endogenous (CD45.2^+^) or exogenous/ALT (CD45.1^+^) **(F)** CD8^+^ T_N_, T_CM_, T_EFF_ proportions between IL-2 and IL-7 ALT in tumor 48-hours following administration. **(G & H)** Comparison of IL-2 and IL-7 ALT phenotype fractions in tumor to ALT pre-administration. **(I & J)** Weight normalized counts at both 3- and 48-hr timepoints of CD8^+^ T_CM_ and CD8^+^ T_EFF_ accumulation in tumor when treating with IL-2 or IL-7 ALT (multiple unpaired t-tests shown). **(K)** Comparison of CD8^+^ T_CM_ and CD8^+^ T_EFF_ presence in tumor following IL-2-ALT or IL-7-ALT. Statistical analyses performed via two-way ANOVA and data presented as mean ± SEM unless otherwise specified. Experimental outline generated using BioRender.com.

Evaluating the phenotypic composition of the CD8^+^ T cells infiltrating tumor, we observed that at 48-hours post ALT administration, the endogenous CD8^+^ T cell phenotypes in tumor were similar, regardless of whether ALT was expanded with IL-2 or IL-7. Phenotypes consisted mainly of effector (T_EFF_) cells (Figure 3E). Regarding administered (exogenous) ALT CD8^+^ T cells, IL-2-ALT in tumor also pre-dominantly consisted of T_EFF_ cells, while IL-7-ALT consisted of a mixed central/effector (T_CM_/T_EFF_) population (Figure 3F). Naïve populations were negligible and did not vary between groups. Given this, we considered that enhanced entry or retention might be more characteristic of the T_CM_ population. However, we noted that the make-up of ALT in tumor was similar to the phenotypic make-up of the IL-2 and IL-7 ALT cell product pre-administration (IL-2-ALT in vitro (pre-administration) vs. in vivo (post-administration) in Figure 3G; IL-7-ALT in Figure 3H). This led us to consider instead whether the increased fraction of CD8^+^ T_CM_ in tumor for the IL-7-ALT group simply reflected the composition of the pre-administration product.

To clarify, we compared accumulation of the CD8^+^ T_CM_ and T_EFF_ subsets between both ALT groups. Both the IL-7-ALT CD8^+^ T_EFF_ and T_CM_ subsets were present in tumor in significantly greater numbers than their IL-2-ALT counterparts (individual subsets for each ALT condition in Figures 3I and 3J, comparison between ALT conditions in Figure 3K, IL-7-ALT vs IL-2-ALT T_CM_ and T_EFF_ p<0.0001****). We concluded that enhanced accumulation of IL-7-ALT CD8^+^ T cells in tumor was more a feature of IL-7 expansion, and not reflective of T_CM_ phenotype.

### Expansion with IL-7 upregulates expression of the pro-migratory integrin VLA-4 on murine CD8^+^ T cells, which is required for enhanced intratumoral accumulation

To better understand the impact of IL-7 on T cell accumulation in tumor, we analyzed the expression of migratory integrins common to all lymphocytes (VLA-4, LFA-1) on exogenous T cells that entered the CNS using flow cytometry (gating strategy in Figure 4A). We observed that both the expression of the migratory integrin VLA-4 and the fraction of VLA-4^Hi^ cells was significantly increased amidst IL-7-ALT CD8^+^ T cells arriving early in tumor compared to IL-2-ALT, though this difference subsided by the 48-hour sampling point (gMFI in Figure 4B, 3-hrs & 48-hrs p<0.01**, CD8^+^ T cell VLA-4^Hi^ % in Figure 4C). Conversely, we saw increased LFA-1 expression on IL-2-ALT CD8^+^ T cells compared to IL-7-ALT at the first sampling timepoint (Figure 4D, p<0.01** LFA-1 gating in Figure S4A). The percentage of CD8^+^ LFA-1^Hi^ T cells was similarly high throughout for both conditions (Figure 4E). The pre-administration fraction of CD8^+^ VLA-4^Hi^ T cells was also higher amidst the IL-7-ALT, while the fraction of pre-administration CD8^+^ LFA-1^Hi^ T cells was similar between groups (CD8^+^ VLA-4^Hi^ T cells in Figure 4F, CD8^+^ LFA-1^Hi^ T cells in Figure 4G, technical repeats for each ALT condition shown).

**Figure 4.**
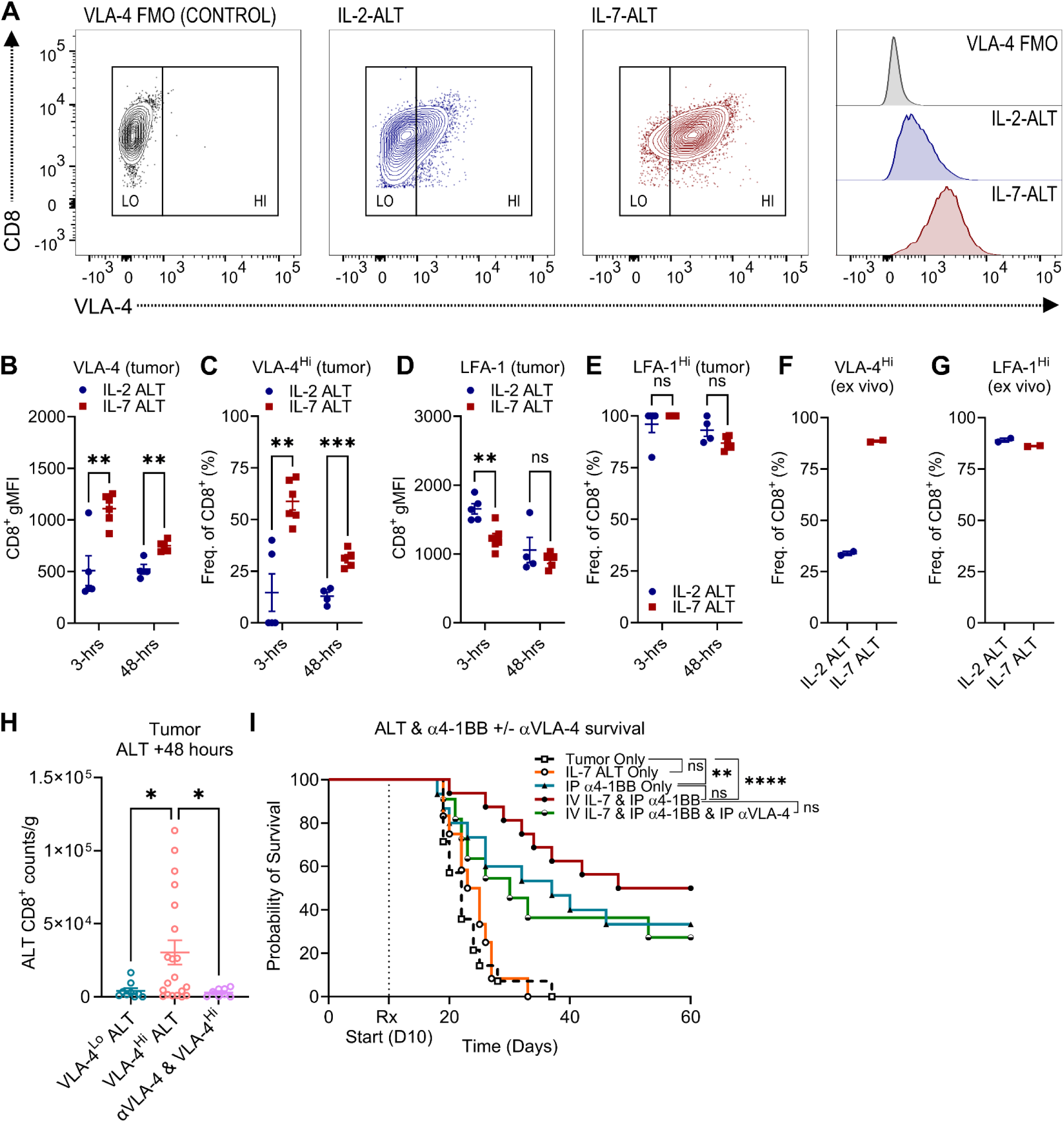
Expansion with IL-7 upregulates expression of the pro-migratory integrin VLA-4 on murine CD8^+^ T cells, which is required for enhanced intra-tumoral accumulation. **(A)** Representative gating strategy shown for VLA-4 expression on CD8^+^ T cells from IL-2 or IL-7 ALT. **(B & C)** CD8^+^ T cell VLA-4 expression (gMFI) and proportion of CD8^+^ VLA-4^Hi^ T cells shown when expanding with IL-2 or IL-7 ALT (experiment outline in Figure 3, n=4-6/group). **(D & E)** CD8^+^ T cell LFA-1 expression (gMFI) and proportion of CD8^+^ LFA-1^Hi^ T cells shown when expanding with IL-2 or IL-7 ALT. **(F & G)** Analysis of the CD8^+^ T cell VLA-4^Hi^ or LFA-1^Hi^ proportion in ALT cellular product at expansion end (2 technical replicates). **(H)** Entry of CD45.1^+^CD8^+^ T cells in tumor following VLA-4^Lo^, VLA-4^Hi^ ALT or VLA-4^Hi^ ALT & αVLA-4 (single-dose 200µg intraperitoneal αVLA-4 antibody (BioXCell) pre-ALT, 3 pooled experiments shown, n=8-20/group, one-way ANOVA shown). **(I)** Evaluation of VLA-4 expression on the endogenous CD8^+^ T cell compartment over time (IL-2 and IL-7 treatment groups pooled, n=9-12/group). Survival comparisons performed via log-rank (Mantel-Cox) chi-squared test. Statistical analyses performed using unpaired T tests and data presented as mean ± SEM unless otherwise specified.

To evaluate the impact of VLA-4 levels on T cell trafficking to tumor, we again used IL-7 to generate an ALT consisting of CD4^+^ and CD8^+^ T cells that had increased VLA-4 expression compared to baseline levels seen when expanding with IL-2 (VLA-4^Hi^ or VLA-4^Lo^ ALT, pooled ALT expression shown in Figure S4B). VLA-4^Hi^ or VLA-4^Lo^ ALT were then separately administered to animals with 15-day established CT2AvIII tumors, either alone or alongside a VLA-4 blocking antibody (BioXCell, 200μg intraperitoneally (IP)). VLA-4^Hi^ ALT CD8^+^ T cells demonstrated significantly enhanced accumulation within tumor compared to VLA-4^Lo^ ALT, accumulation that was abrogated by co-administration of αVLA-4 (VLA-4^Hi^ vs VLA-4^Lo^, p<0.05*; VLA-4^Hi^ vs αVLA-4 & VLA-4^Hi^, p<0.05*, 3 pooled experiments shown, Figure 4H). Administration of αVLA-4 was also found to reduce the therapeutic efficacy of combination IL-7-ALT + α4-1BB seen in the CT2AvIII model. Here, we observed that IP αVLA-4 reduced survival in the combination therapy group from 54 to 30 days, similar to monotherapy groups, though these curves were not significantly different (IP α4-1BB only: 37 days, 3 pooled experiments shown, Figure 4I). Based on these findings, we concluded that: (1) IL-7 exposure increases the expression of VLA-4 on CD8^+^ T cells and (2) VLA-4 expression and signaling are necessary for the enhanced accumulation of IL-7 ALT CD8^+^ T cells in glioma.

### Intratumoral lymphocytic VLA-4 and endothelial/pericytic VCAM-1 expression increase over time in glioma

While evaluating ALT VLA-4 expression, we noted that VLA-4 levels were consistently elevated over time on endogenous CD8^+^ T cells in tumor (Figure 5A). Noting the role of VLA-4 in guiding T cells towards inflamed tissue (38), we first examined whether VLA-4^Hi^ T cells were unique to the tumor environment. We performed flow cytometry on various tissues (brain, spleen, bone marrow, lungs, and blood) in mice with established CT-2AvIII tumors or sham-injected controls (experimental outline in Figure 5B, gating strategy in Figure S5A). Animals did not receive ALT, allowing for evaluation of endogenous T cell VLA-4 expression without external influence. We again observed a significant increase to the VLA-4^Hi^ fraction amidst CD8^+^ T cells, as well as to weight-adjusted VLA-4^Hi^ CD8^+^ T cell counts in brain tumor tissue compared to all other compartments, as well as to brain from sham controls (fractions in Figure 5C, comparisons in Figure 5D, p<0.0001****, weight adjusted counts vs sham controls in Figure S5B, p<0.01**). VLA-4^Hi^ CD8^+^ T cells in tumor were overwhelmingly T_EFF_ cells (Figure S5C). Outside of the tumor, VLA-4^Hi^ CD8^+^ T cells were most commonly found in the bone marrow (Figure 5, C and D).

**Figure 5.**
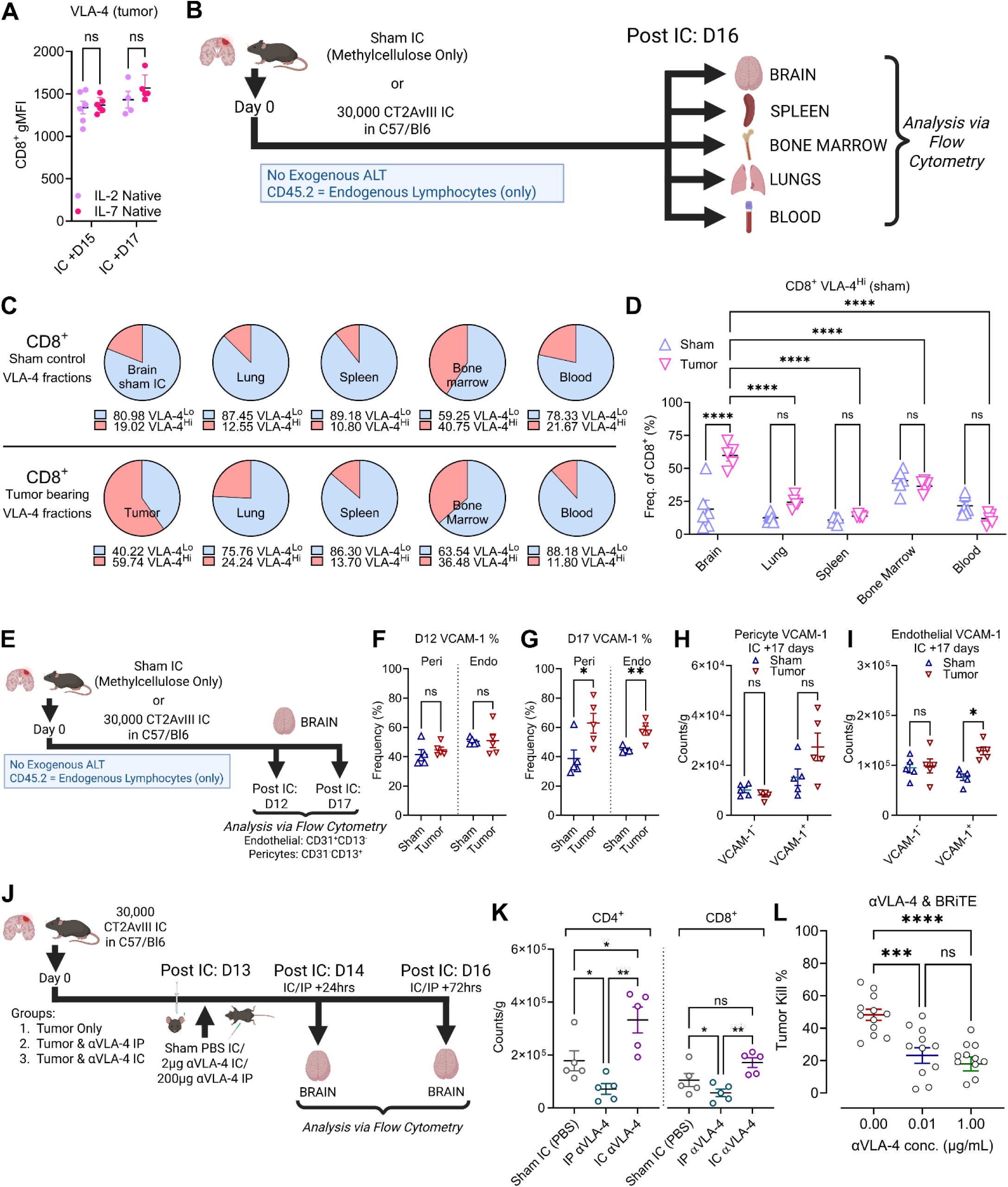
Lymphocytic VLA-4 & endothelial/pericytic VCAM-1 expression increases over time on native T cells in murine glioma. **(A)** Comparison of CD8^+^ T cell VLA-4 gMFI expression on endogenous cells over time from prior experiment (n=4-6/group). **(B)** Study overview to assess VLA-4 expression across compartments. **(C & D)** Evaluation of VLA-4^Lo^ and VLA-4^Hi^ fractions across different compartments. Comparisons in **D** via two-way ANOVA (n=5-6/group). **(E)** Study overview to assess endothelial VCAM-1 in the CNS. No ALT was used. Tumor hemispheres were collected, and endothelial cells (CD31^+^CD13^-^)/pericytes (CD31^-^CD13^+^) were analyzed (n=5/group). **(F & G)** Frequency of VCAM-1^+^ pericytes/endothelial cells in tumor at D12 and D17 following tumor/sham injection. **(H & I)** Counts of VCAM-1^-^/VCAM-1^+^ pericytes/endothelial cells in tumor compared to sham controls at D17 following intracranial injections. **(J)** Study overview to evaluate intraperitoneal/intracranial VLA-4 blockade (n=5/group). **(K)** Intracranial CD4^+^ & CD8^+^ T cell counts 72-hours following administration of intraperitoneal/intracranial αVLA-4 or sham controls. Comparisons via one-way ANOVA. **(L)** In vitro cytotoxicity with CT2AvIII, T cells, αVLA-4 and BRiTE at EC_50_ concentration of 0.01µg/mL. n=12/dose level. Comparisons via one-way ANOVA. Statistical analyses performed using unpaired T tests and data presented as mean ± SEM unless otherwise specified. Experimental outlines generated using BioRender.com.

As glioblastoma is characterized by high levels of microvascular proliferation (39), we next evaluated if endothelial/pericytic VCAM-1 (VLA-4’s ligand (40)) was also increased in tumor. Recent studies of T cell recruitment in other intracranial tumor models (i.e., melanoma) found negligible VCAM-1^+^ endothelial or pericyte populations (41). To determine if VCAM-1^+^ cells were present in the context of glioma, we collected brain tissue from mice with established intracranial CT2AvIII glioma or sham-controls. Both endothelial (CD31^+^CD13^-^) and pericyte (CD31^-^CD13^+^) populations were analyzed via flow cytometry (outline in Figure 5E, gating in Figure S5D). Here, we observed a similar frequency of expression of VCAM-1 on endothelial cells and pericytes on Day 12 of tumor growth (∼45%^+^), with similar levels also seen across tumor and sham (Figure 5F). However, as tumors progressed further, increasing proportions and counts of VCAM-1^+^ endothelial cells and pericytes were observed in tumor compared to sham (Figure 5G, pericytes p<0.05*, endothelial cells p<0.01**, D17 weight adjusted counts in Figure 5, H and I respectively).

Given this, we questioned whether intratumoral T cell accumulation might be enhanced by blocking the VLA-4-VCAM-1 axis, and whether results would differ if blockade was administered systemically vs. intracranially. To evaluate, we implanted mice with CT2AvIII gliomas, and 13 days later administered αVLA-4 either intraperitoneally or intracranially. As free αVLA-4 antibody levels are 100-fold reduced in cerebro-spinal fluid (CSF) compared to serum following systemic infusion (42), we selected dose levels of 200 µg αVLA-4 intraperitoneally and 2 µg intracranially to achieve approximately equivalent exposure (experiment outline in Figure 5J). First, we evaluated the impact of both intraperitoneal and intracranial αVLA-4 on intratumoral CD8^+^ T cell VLA-4 expression and found that both suppressed VLA-4 levels on T cells, though this was only sustained when administering intraperitoneally (Figure S5E). Next, we evaluated intracranial counts of endogenous CD4^+^ and CD8^+^ T cell populations 24- and 72-hrs post αVLA-4 administration. Across treatment groups, no differences were seen at the 24-hour measurement point (Figure S5F). By 72-hours, however, intraperitoneal αVLA-4 had elicited reduced counts of both CD4^+^ and CD8^+^ T cells in tumors, even compared to non-tumor-bearing controls (24-hr intratumoral counts in Figure S5F, 72-hr counts in Figure 5K, intraperitoneal αVLA-4 vs sham, CD4^+^ and CD8^+^ T cells, p<0.05*). Conversely, intracranial αVLA-4 produced increases in accumulation of both populations within tumors by 72-hrs post administration, though this was only significant for CD4^+^ T cells compared to sham controls (Figure 5K, intracranial αVLA-4 vs sham, CD4^+^ T cells p<0.05*, CD8^+^ T cells p=0.068).

We therefore further investigated the therapeutic potential for intracranial VLA-4 blockade. While increasing intratumoral T cell counts remains a central goal, VLA-4 is involved in immune synapse formation and T cell activation (43), making the net impact of its blockade unclear. To evaluate the effects of VLA-4 blockade on T cell activity, we performed in vitro co-culture cytotoxicity assays with CT2AvIII, T cells, and EGFRvIII-BRiTE. αVLA-4 antibody was added at escalating amounts to BRiTE at its EC_50_ dose (0.01µg/mL, dose-finding data in Figure S5G). The addition of αVLA-4 indeed inhibited the T cell anti-tumor cytotoxicity otherwise mediated by BRiTE (Figure 5L). Thus, while intracranial αVLA-4 may enhance intratumoral T cell accumulation, there appears to be a negative impact on anti-tumor cytotoxicity.

### IL-7 expansion of hPBMCs from both healthy volunteers and patients with glioblastoma upregulates lymphocyte VLA-4 expression

We therefore returned to further investigate our initial IL-7-ALT method for enhancing T cell numbers in tumor. To determine the clinical translatability of our approach, we sought to determine whether a) leukapheresis products from patients with glioblastoma would respond / expand to the same extent as those from healthy controls, and b) similar changes to VLA-4 expression and T cell migratory potential would occur when expanding human T cells in the presence of IL-7.

To permit relevant comparisons, we obtained leukapheresis products from healthy volunteers and patients with glioblastoma who had given written consent to provide PBMCs for the purposes of T cell expansion. An overview of donor demographics and pathology is shown in Table 1. Median ages across groups were similar (controls 54.3 yrs (range 43.3– 55.4) vs glioblastoma 58.7 yrs (range 37.4–63.3)) with a 2:1 M:F ratio in both groups. Of the glioblastoma samples, 5 of 6 were collected from patients who had previously received dexamethasone.

**Table 1.** Demographics of hPBMC samples taken from healthy volunteers and glioblastoma patients.

| Characteristics | Control<br>N = 5 | Glioblastoma<br>N = 6 |
| --- | --- | --- |
| **Age (yr)** |  |  |
| Median | 54.3 (43.3–55.4) | 58.7 (37.4–63.3) |
| **Sex** |  |  |
| Male | 2/5 (40%) | 4/6 (66%) |
| Female | 1/5 (20%) | 2/6 (33%) |
| Unknown | 2/5 (40%) | 0 |
| **Ethnicity** |  |  |
| American Indian/Alaska Native | 0 | 0 |
| Black or African American | 0 | 0 |
| Hispanic or Latino | 0 | 0 |
| Native Hawaiian or other Pacific Islander | 0 | 0 |
| Caucasian | 3/5 (60%) | 6/6 (100%) |
| Unknown | 2/5 (40%) | 0 |
| **Diagnosis** |  |  |
| IDHwt* glioblastoma | NA | 6/6 (100%) |
| Newly diagnosed | NA | 4/6 (66.7%) |
| Recurrent | NA | 2/6 (33.3%) |
| **Steroid Status** |  |  |
| Naïve | NA | 1/6 (16.7%) |
| Experienced | NA | 5/6 (83.3%) |

\*IDHwt: Isocitrate Dehydrogenase wild-type

Samples from control and glioblastoma groups were thawed and activated via αCD3/αCD8 stimulation. Cultures were maintained for 14 days in media supplemented with 300 IU/mL IL-2 (based on prior protocols (44)) or 20 ng/mL IL-7. After expansion, cell counts, phenotypes, activation levels and VLA-4 expression were evaluated (flow gating and schematic in Figure S6, A and B). While both IL-7 and IL-2 were capable of independently supporting T cell proliferation (Figure 6A), IL-2 produced greater T cell yields for both control and GBM leukapheresis samples (Figure S6C). Perhaps surprisingly, no significant difference between control and glioblastoma T cell expansion rates were observed. The final expanded product consisted of ∼95% CD3^+^ T cells across all groups (Figure S6D). IL-7 expansion skewed significantly more towards CD4^+^ T cells (∼60:40 ratio of CD4^+^:CD8^+^ T cells), whereas IL-2 skewed more towards CD8^+^ T cells (CD4^+^:CD8^+^ T cells ∼ 33:66, Figure 6, B-D). On more specific phenotyping of memory and effector subsets, we observed that T_EFF_ (CD45RO^-^ CD45RA^+^CD95^+^CCR7^-^) and T Effector Memory (T_EM,_ CD45RO^+^CD45RA^-^CD62L^-^CCR7^-^) fractions were similar across the IL-2 and IL-7 expansion groups (T_EFF_ in Figure S6E, T_EM_ in Figure S6F, all subsets in Figure 6E). IL-7 did trend toward increasing the stem cell memory fraction (T_SCM,_ CD45RO^-^CD45RA^+^CD95^+^CCR7^+^) compared to IL-2, though changes were non-significant (Figure 6F).

**Figure 6.**
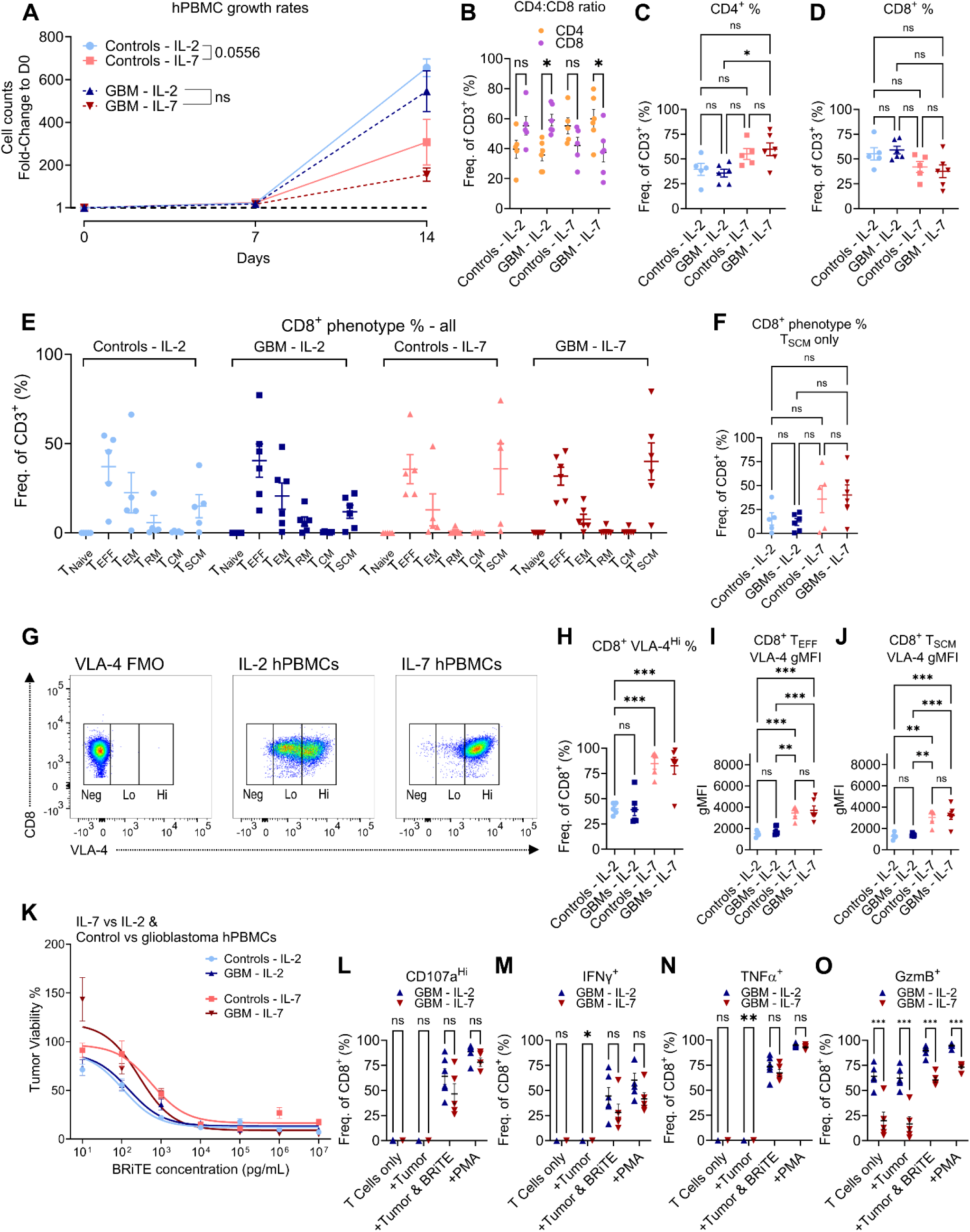
IL-7 expansion of hPBMCs from both healthy volunteers and patients with glioblastoma upregulates lymphocytic VLA-4 expression. **(A)** hPBMC growth rates for IL-2 vs IL-7 co-culture from both glioblastoma and control volunteer leukaphereses. (n=5-6/group). **(B-D)** CD4^+^:CD8^+^ T cell ratios and comparisons for IL-2 vs IL-7 for control and glioblastoma samples at expansion end. **C** & **D**. Comparisons in B via two-way ANOVA and in C & D via one-way ANOVA. **(E)** Overview of CD8^+^ T cell phenotype fractions at expansion end: Naïve (T_N_), Effector (T_EFF_), Effector Memory (T_EM_), Resident Memory (T_RM_), Central Memory (T_CM_), Stem Cell Memory (T_SCM_). **(F)** Fraction of CD8^+^ T_SCM_ at expansion end across groups. **(G-J)** Gating strategy for VLA-4^Hi^ fraction and proportion of VLA-4^Hi^ in all CD8^+^ T cells (**H**) as well as CD8^+^ T_EFF_ (**I**) and CD8^+^ T_SCM_ cells (**J**) at expansion end. **(K)** Cytotoxicity assay with tumor (U87vIII) co-cultured with IL-2 or IL-7 expanded donor/glioblastoma hPBMCs & BRiTE. **(L-O)** Degranulation assays assessing CD107^Hi^, GzmB^+^, IFNγ^+^, TNFα^+^ in glioblastoma CD8^+^ T cells expanded with IL-2 and IL-7. Comparisons via multiple unpaired t tests (n=5-6/group). Statistical analyses performed using one-way ANOVA and data presented as mean ± SEM unless specified.

We next evaluated whether IL-7 increased VLA-4 expression on human T cells, as we had seen in our mouse models. We observed a biphasic distribution in VLA-4^+^ cells and classed lymphocytes as VLA-4^Neg^, VLA-4^Lo^ or VLA-4^Hi^ (gating in Figure 6G). For both control and glioblastoma samples, IL-7 significantly increased the VLA-4^Hi^ fraction over that seen with IL-2 (Figure 6H, p<0.001***, CD4^+^ and CD8^+^ T cell gMFI shown in Figure S6, G & H). Significant upregulation of VLA-4 expression was observed on CD8^+^ T_EFF_ and CD8^+^ T_SCM_ subsets from both groups (Figure 6, I and J, p<0.01** for healthy controls and p<0.001*** for glioblastoma patients). Changes to CD8^+^ T_EM_ VLA-4 expression were non-significant (Figure S6I).

Finally, we performed in vitro functional assays. These included cytotoxicity assays using our end-expansion product, cultured alongside BRiTE and EGFRvIII^+^ positive tumor (U87vIII), as described above. Similar dose-dependent tumor killing was observed across all conditions above BRiTE concentrations of 10^3^ pg/mL (Figure 6K). We then undertook in vitro stimulation assays of sorted CD8^+^ T cells (cytokine gating strategy in Figure S7A, sort gating in Figure S7B). Sorted CD8^+^ T cells were expanded with IL-2 or IL-7 and co-cultured with tumor only (U87vIII); tumor + BRiTE; or phorbol myristate acetate (PMA)/ionomycin. T cells not exposed to tumor, BRiTE or PMA were also analyzed to assess background activity. We observed similar results for IL-2- and IL-7-expanded glioblastoma patient CD8^+^ T cells in terms of CD107a^Hi^ degranulation, interferon-gamma (IFN-γ), and tumor-necrosis factor-alpha (TNF-α) production (Figure 6, L-N). While we did observe increased frequencies of Granzyme B^+^ (GzmB^+^) T cells in the IL-2-expanded group compared to those expanded with IL-7, we also noted an increased baseline frequency of GzmB^+^ T cells in the unstimulated / IL-2-expanded and tumor only control groups (Figure 6O). Ultimately, then, IL-7 expansion enhanced CD8^+^ T cell VLA-4 expression amongst hPBMC samples in a manner similar to that seen with murine T cells, while having a similar impact on T cell function to IL-2 exposure.

### IL-7 increases transcription of genes in T cells associated with enhanced migratory function (*CD9*) and protection against sequestration (*S1PR1)*

To evaluate IL-7’s effect on migratory integrin expression, we performed bulk RNA sequencing using the same glioblastoma hPBMC samples. CD8^+^ T cells were sorted from naïve (pre-expansion), IL-2-expanded, or IL-7-expanded products. Principal component analysis revealed that expansion with IL-7 vs. IL-2 resulted in distinct CD8^+^ T cell transcriptional fates (Figure 7, A-C). Deeper interrogation of these fates via analysis of differential gene expression revealed that IL-7 significantly increases the transcription of genes involved in T cell migration and trafficking compared to IL-2 (*EPHA4* (adjusted p (p_adj_)= 9.39E-27(45))*, ITGA4* (VLA-4, p_adj_=0.043) and *ITGA6 (*VLA-6, p_adj_=9.64E-08 (46))). Genes involved in stem-like/memory T cell development were also significantly increased by IL-7 exposure (*TCF7* which encodes for TCF1, p_adj_=1.60E-06 (47), *FOXO1,* p_adj_=7.52E-11 and *LEF1*, p_adj_=2.41E-10). IL-2-expanded CD8^+^ T cells demonstrated increased transcription of T cell markers associated with effector activity (*CD69,* p_adj_=2.48E-34*, GZMB,* p_adj_=2.16E-18) and exhaustion (*CTLA-4,* p_adj_=1.55E-33, *TOX* p_adj_=0.00068, *HAVCR2* (TIM-3) p_adj_=0.00018). Volcano plots for all three comparisons are shown in Figure S8, A and B (naïve vs IL-2 or IL-7, respectively), and Figure 7D (IL-2-vs IL-7-expanded). Analysis of raw transcripts per million (TPM) also found that IL-7 yielded the highest TPM for *ITGA4* (VLA-4), but this was not significantly increased compared to the IL-2 cohort (Figure S8C).

**Figure 7.**
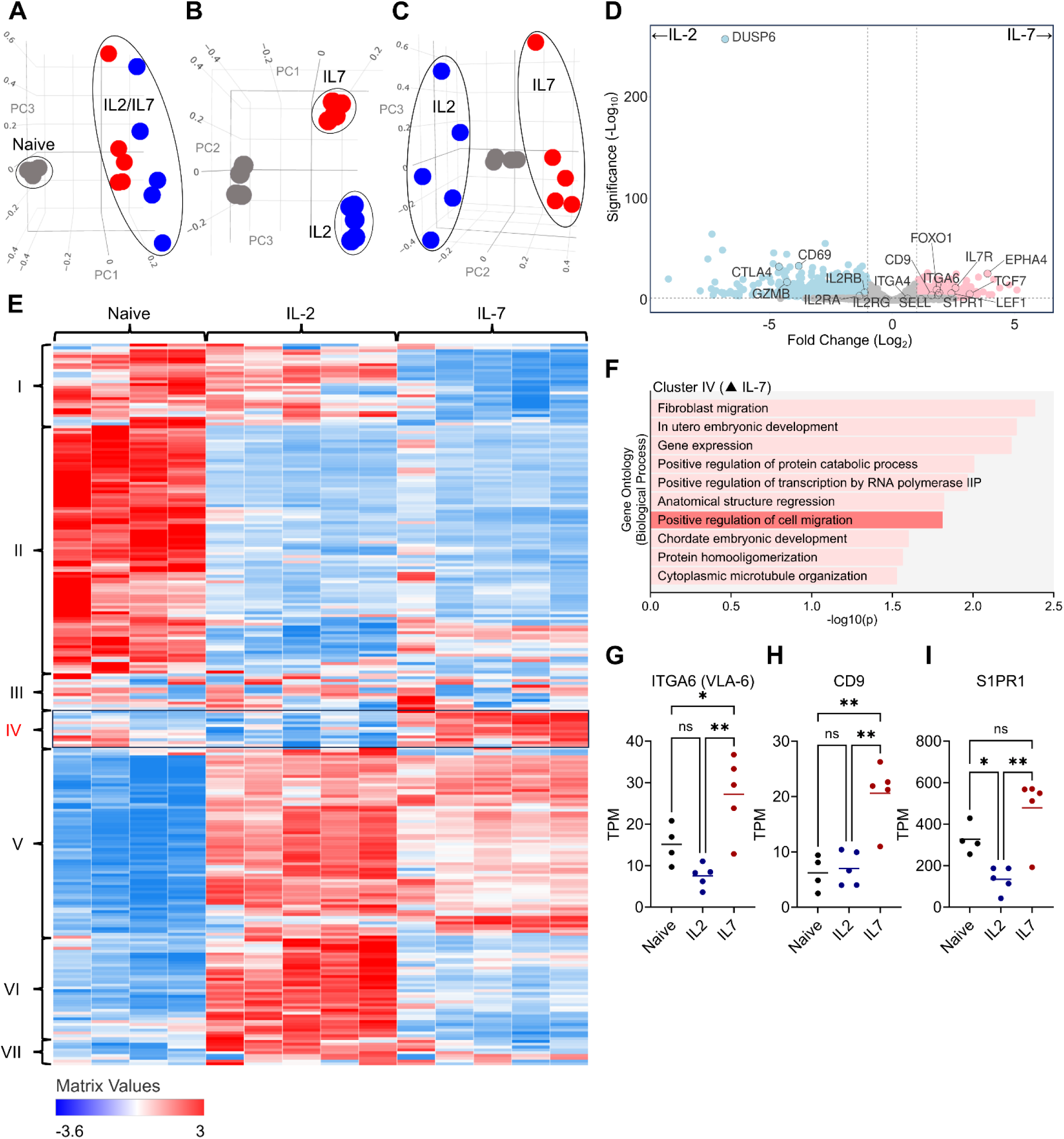
IL-7 upregulates transcription of genes associated with enhanced migratory integrin expression (*CD9*) and protection against tumor-induced sequestration (*S1PR1*) **(A-C)** Principal component analysis of naïve (grey), IL-2 expanded (blue) and IL-7 expanded (red) CD8^+^ T cells (n=4-5/group for all graphs). **(D)** Volcano plot comparing differential gene expression between IL-2 and IL-7 expanded CD8^+^ T cells. **(E)** Hierarchical cluster heatmaps of the naïve, IL-2 and IL-7 groups with seven unique clusters identified (I-VII). **(F)** Gene Ontology biological processes analysis of genes expressed in cluster IV (upregulated with IL-7, *EPHA4, CD9, ITGA6, S1PR1* etc.) **(G-I)** Comparisons of raw transcripts per million (TPM) across groups for *ITGA6, CD9* and *S1PR1* (comparisons via one-way ANOVA). PCA analysis was generated using BioJupies (66). Hierarchical cluster heatmaps were generated using clustergrammer (Maayan lab, (67)). Gene Ontology (GO) biological processes analysis was performed using the NIH DAVID database (53). Volcano plots were generated using VolcaNoseR (69). Comparisons via one-way ANOVA and data presented as mean ± SEM unless otherwise specified.

Differential expression analysis also revealed a highly significant outlier when expanding with IL-2 vs IL-7. Transcription of Dual-Specificity Phosphatase 6 (*DUSP6*) was increased 6.8x when expanding with IL-2 vs IL-7 (Figure 7D, p= 7.85E-258). DUSP6 has been shown to negatively regulate the extracellular signal-regulated kinase (ERK1/2) pathway (48–50), which is activated by IL-2 (51). Curious as to the impact of DUSP6, we performed a repeat 2-week expansion where hPBMCs were expanded with IL-2 alone, or alongside BCI, a small molecule inhibitor of DUSP6 (52). We found that while BCI did not increase CD8^+^ T cell VLA-4 expression, it did increase the fraction of CD8^+^ T_SCM_ cells (CD8^+^ T cell VLA-4 gMFI in Figure S8D, CD8^+^ T_SCM_ fraction in Figure S8E, technical repeats). We surmised that DUSP6 may affect CD8^+^ T cell differentiation, but not VLA-4 expression.

Noting that transcription of another migratory integrin (VLA-6) was enhanced with IL-7, we considered whether broader changes in integrin expression might be occurring. We performed hierarchical cluster heatmapping of genes that were differentially expressed across the naïve, IL-2 and IL-7 groups. This identified seven clusters (I-VII, Figure 7E). Clusters I and VI were specifically upregulated in the IL-2-expanded samples, while cluster V was upregulated for both. Cluster IV was the only gene set specifically upregulated within IL-7-expanded samples. To interrogate the biological processes that these clusters represented, we performed Gene Ontology (GO) analysis using the NIH DAVID database (53). For clusters I and VI (upregulated in IL-2), we identified biological processes associated with an inflammatory adaptive immune response and positive regulation of T cell activation (Figure S8, F and G). These included genes related to cytotoxic T cell activity (*GZMA, GZMB, IFNγ, FASLG, CD69*) as well as markers of T cell exhaustion (*CTLA-4, TOX*). For cluster V (upregulated in both), increased transcription of genes associated with cell division were observed (e.g., *MKI67* (*54*), Figure S8H).

GO analysis of cluster IV (upregulated in IL-7) identified biological processes associated with positive regulation of cell migration (Figure 7F). These included genes encoding for *ITGA6* (VLA-6, TPM comparisons across groups shown in Figure 7G) as well as *CD9*, a protein that has been associated with increased expression of multiple migratory integrins, including VLA-6 and VLA-4 (55, 56). Within our data, *CD9* TPM was significantly and specifically increased within IL-7 expanded hPBMCs compared to naïve or IL-2 expanded cohorts (p<0.01**, Figure 7H). Interestingly, we also identified *S1PR1* (encoding the S1P1 receptor) within cluster IV. Our group has previously reported that loss of S1P1 mediates tumor-induced T cell sequestration in bone marrow (5) and that stabilization of S1P1 on the surface of T cells makes them resistant to sequestration (5). Here, we find that IL-2 significantly reduced *S1PR1* TPM compared to naïve controls (p<0.05*), while IL-7 significantly increased *S1PR1* TPM (p<0.001***, Figure 7I). We concluded that IL-7 upregulates genes associated with migratory integrin expression, while also increasing *S1PR1* transcription.

## Discussion

For T cell-activating therapies to be effective against tumors of the CNS, they must be able to encounter functional immune cells within the intracranial compartment. Although immune cells can cross the BBB (28), primary CNS malignancies are capable of “driving” T cells away by inducing loss of lymphocytic S1P1, resulting in T cell sequestration within bone marrow (5, 57).

Our results demonstrate that non-specific CD8^+^ T cells expanded with IL-7 can accumulate in tumor tissue, despite tumor-induced T cell sequestration. Further, we demonstrate that the administration of IL-7-ALT can synergize with multiple T cell-activating therapies against established intracranial glioma. Campian et al reported a related phenomenon, whereby peripheral administration of long-acting recombinant IL-7 enhanced cytotoxic CD8^+^ T cell numbers systemically and in tumor for similar animal glioma models (13). Another recent study by Lee et al. reports synergism between recombinant IL-7 administration and bi-specific T cell engager therapy in solid tumor models (27). Interestingly, they report that IL-7 encourages recruitment of non-exhausted polyclonal bystander CD8^+^ TILs to tumors. Though these bystander TILs do not have direct anti-tumor activity themselves, they can be redirected by T cell engagers to become tumoricidal. Our study adds to this body of literature, finding that IL-7 enhances expression of migratory integrins like VLA-4, and increases transcription of genes involved in resistance to sequestration (*S1PR1*).

IL-7’s effect on upregulating VLA-4 has been previously reported in vitro (58), and peripheral administration of recombinant human IL-7 (rhIL-7) has been observed to increase VLA-4 and T cell trafficking in murine models of sepsis (59). In our study, RNA sequencing of CD8^+^ T cells from glioblastoma patients similarly identified that transcription of VLA-4 was greatest when expanding with IL-7. Interestingly, while only IL-7 significantly increased CD8^+^ VLA-4 transcription levels compared to pre-expansion lymphocytes, no significant difference in VLA-4 TPM was observed between IL-2 and IL-7 expanded cells. However, evaluation of genes uniquely upregulated by IL-7 did identify transmembrane proteins *(CD9)* associated with multiple VLA integrins (55, 56). In keeping with this finding, we observed significantly increased transcription of *ITGA6* (VLA-6) when expanding with IL-7 compared to IL-2. Though our study found that VLA-4 was necessary for enhanced CD8^+^ T cell accumulation, it is possible that IL-7 increases the expression of multiple pro-infiltrative integrins. These could further enhance the ability of CD8^+^ T cells to enter and accumulate in intracranial glioma.

As mentioned, other genes uniquely upregulated by IL-7 included the g-protein-coupled receptor *S1PR1*. We also have previously identified that S1P1 loss mediates T cell sequestration (5). Accordingly, mice with genetically stabilized S1P1 expression in T cells prove sequestration-resistant and experience prolonged survival to immunotherapy in the face of glioma (5). Our findings align with this prior work. Notably, we also identify that IL-2 exposure decreases the transcription of *S1PR1*. This in turn may underlie some of the failure of IL-2-expanded cellular products to accumulate in intracranial tumors, though further confirmatory work is required.

We also report here increasing lymphocyte VLA-4 expression and increasing numbers of VCAM-1^+^ endothelial cells/pericytes in the context of glioma over the course of tumor progression. Given that recent studies have reported low fractions of VCAM-1^+^ cells in other intracranial malignancies (melanoma (41)), this finding suggests that T cell recruitment and BBB trafficking dynamics may vary between cancer types. To identify disease-specific approaches for enhancing T cell accumulation in tumors of the CNS, further evaluation of VCAM-1^+^ expression across multiple tumor types is warranted. Nevertheless, our findings suggest that for intracranial glioma, VLA-4^Hi^ lymphocytes may be able to encounter VCAM-1 and accumulate in tumor.

To better reflect the capacity for clinical translation, we confirmed that IL-7 exposure upregulated VLA-4 expression on hPBMCs from both healthy donors and patients with glioblastoma. While we observed reduced yields and less CD8^+^ T cell skewing when expanding hPBMCs with IL-7, we did find that IL-7-cultured hCD8^+^ T cells exhibited similar in vitro functionality to those being IL-2-cultured. Further optimization could include sorting infiltrative (VLA-4^Hi^) CD8^+^ T cells pre-administration. This might also permit administering a reduced dose to patients, or one with more efficient tumor uptake. In our study, the percentage of the administered ALT dose which entered the brain was small, in keeping with other studies evaluating IV CAR-T therapy against murine CNS malignancy models (<1%, lymphoma (60)). Though our approach did not induce overt toxicity/demyelination in brain or spinal cord when combining IL-7-ALT with αPD-1, we did observe a few scattered foci of hepatic cellular apoptosis/necrosis associated with lymphocytic infiltrate. An optimized VLA-4^Hi^ population could reduce the ALT dose required for efficacy, and limit non-CNS organ infiltration.

A key limitation of our study is that we only evaluated the cytokines IL-2 and IL-7, as we have previously established that they can independently support T cell expansion (22). Similarly, we used doses of IL-2 and IL-7 that are known to expand T cells from both mice and humans. However, cytokines such as IL-15, IL-18, IL-21 and others have been used in varying doses to supplement lymphocyte expansion (61). We also note that the IL-7 receptor is downregulated after sustained exposure, and intermittent cycling with IL-7 may enhance the cytokine’s effects (62). Our study is confined to evaluating these two cytokines at the specified concentrations and exposure schedules. However, given the vast number of possible permutations, determining the optimal combination, concentration, and timing for cytokine supplementation is beyond the scope of this manuscript and may require a continuous process-development effort.

In conclusion, we find that expansion of autologous T lymphocytes with IL-7 enhances the ability of CD8^+^ T cells to accumulate within intracranial glioma, even in the setting of tumor-imposed T cell sequestration. Combination IL-7-ALT and T cell-centric immunotherapies increase survival in multiple models of glioma. Analysis of migratory integrins on lymphocytes finds that IL-7 increases CD8^+^ T cell expression of VLA-4. RNAseq analysis finds that IL-7 increases transcription of genes associated with increased migratory integrin expression *(CD9)* as well as *S1PR1*, a G protein-coupled receptor whose stable expression on lymphocytes protects against their sequestration. These findings will be used to inform future clinical trials, where ALT pre-treatment will be combined with T cell-activating therapies targeting glioblastoma (e.g., BRiTE, NCT04903795).

## Materials and Methods

### Sex as a biological variable

For pre-clinical in vivo studies, we exclusively used females as CT-2A was established in female mice (63) and superior engraftment of human hematopoietic stem cells has been observed in female NOD-*scid* gamma (NSG) mice (64). Evaluation of clinical samples from volunteers and glioblastoma patients used hPBMCs from males and females (**Table 1**). Further demographic information in terms of reporting race and ethnicity were based on classifications within the electronic medical records system where available and categorized according to NIH policy (notice number: NOT-OD-15-089).

### Mice

All mouse strains used, and housing conditions are described within the Supplemental Methods.

### Cell lines

C57BL/6 syngeneic CT2A was originally provided by Robert L. Martuza (Massachusetts General Hospital, Boston, MA, USA). Generation of stably transfected sublines was performed in-house and used in previous studies (1). Similarly, U87MG was obtained from the American Type Culture Collection (ATCC, cat #HTB-14) and transfected in-house to stably express EGFRvIII. GL261 was obtained from the NCI (National Cancer Institute, Frederick, Maryland, USA). Their generation and usage has also been described in prior studies by our group (2). All cell lines are authenticated and confirmed to be contaminant free via IDEXX Laboratories. CT2A, U87MG and GL261 lines were maintained in culture using complete DMEM (Gibco, 11995-065, 10% FBS) and passaged using 0.05% Trypsin, EDTA (Gibco, 25300-054).

### Murine lymphocyte culturing

Expansion processes for murine CD3^+^ cells are described within the Supplemental Methods.

### Leukapheresis and human PBMC culturing

Human PBMC collection, processing and expansion are described within the Supplemental Methods.

### Tumor inoculation

All tumor studies in this report placed tumors intracranially in mice and followed protocols described previously (65), which are described within the Supplemental Methods.

### In vivo adoptive lymphocyte transfer and antibody administration

For adoptive lymphocyte transfer in this study, cells that had been expanded as described in the murine lymphocyte culturing section were administered as described within the Supplemental Methods.

### Tissue processing and flow cytometry

Procedures for tissue processing and flow cytometry are based on our previously published protocols (4) and are described within the Supplemental Methods.

### Cytotoxicity

Tumor cells were labelled with a viability dye before co-culture. The process is described within the Supplemental Methods.

### Immune functional assays

Immune functional assays were performed with flow-sorted human CD8^+^ T cells and is described within the Supplemental Methods.

### Immunohistochemistry & toxicity

Toxicity studies were conducted in C57/Bl6 mice. Processing procedures are described within the Supplemental Methods.

### RNA sequencing assays and analysis

Flow-sorted CD8^+^ T cells were snap frozen in cell pellets and analyzed via bulkRNA sequencing at Azenta life sciences. Analysis is described within the Supplemental Methods. PCA analysis was generated using BioJupies (66). Hierarchical cluster heatmaps were generated via the same method and using clustergrammer (Maayan lab, (67)). Gene Ontology (GO) biological processes analysis was performed using the NIH database for annotation, visualization and integrated discovery dashboard (DAVID (53)). Volcano plots were generated based on tests for differential expression (Wald test used to generate p-values and log2 fold changes) created with DESeq2 (68) and visualized using VolcaNoseR (69).

### Statistics and reproducibility

Experimental results are presented as mean ± SEM unless otherwise stated. Statistical tests for all studies were completed using GraphPad v.10.3.1 (Prism). For comparisons in a single graph, 2-tailed Student’s t test or nonparametric Mann-Whitney U test was used, and for multiple comparisons, ANOVA (one-way or two-way) with Holm–Šidák correction for multiple comparisons. Asterisks represent the significance level of any difference (\**p*<0.05, \*\**p*≤0.01, \*\*\**p*<0.001, \*\*\*\**p*<0.0001, *p*>0.05 not significant). Sample sizes were selected due to practical considerations, with no formal power calculations performed. Efficacy studies following animals for survival were assessed for significance using a log-rank (Mantel-Cox) chi-squared test. Independent study results were pooled if the effect of replication did not cause significant variation as assessed by two-way ANOVA. All animals were randomized within genotype prior to treatment following tumor implantation. Survival was monitored with the assistance of technicians from the Duke Division of Laboratory Animal Resources (DLAR) who were blinded to study groups and followed animals to endpoint.

### Study approval

All animal experiments were performed under Duke University IACUC experimental protocol (ID: A163-21-08). Stored anonymized human samples used were collected from glioblastoma patients or volunteers who provided written consent to undergo leukapheresis for T cell expansion for research (glioblastoma donors: Duke IRB #Pro00069444; Duke IRB #Pro00083828; healthy donors: Duke IRB #Pro00009403; approval for secondary analysis: Duke IRB #Pro00117663).

### Graphical illustrations

The graphical abstract and experimental outlines in Figs. 1, 2, 3, 5, 6, S1, S3 and S7 were created with BioRender.com and exported under a paid license.

## Data Availability

Values for underlying data for all figures are available in the supporting data values file. RNAseq data (processed TPM counts for individual anonymized samples) has been deposited on the NCBI Gene Expression Omnibus (GEO) repository under accession number GSE288436, available at https://www.ncbi.nlm.nih.gov/geo/query/acc.cgi?acc=GSE288436. Institutional certification for deposition of genomic summary results was granted by Duke’s IRB review board.

## Author Contributions

Design, acquisition, analysis of data, and writing of manuscript: KS with assistance from KMH, SLC, PN, YZ, EM, COR, EEB, BJP, SEW, PKN, GEA, BHS, KA, JHS, MK and PEF. Overall supervision throughout by JHS, MK and PEF.

## Acknowledgements

The authors have been supported by the following grants to Duke University: P50-CA190991 (SPORE) (JHS, MK), P01-CA225622 (JHS, MK), U01-NS090284 (JHS), R01-NS099463 (JHS), R01-CA175517 (JHS), and R01-CA235612 (JHS, MK) as well as a Structured Research Agreement with Adaptin Bio (KS, MK). The authors also wish to express their gratitude to Dr. Rebecca Bacon and Dr. Ivan Spasojevic at Duke University Medical Center for their assistance with toxicity studies.

