## Supplemental methods, figures and tables for "IL-7-mediated expansion of autologous lymphocytes increases CD8^+^ VLA-4 expression and accumulation in glioblastoma models"

#### Mice

Six-to ten-week-old C57BL/6J were purchased from Charles River (cat #556). CD45.1 (B6.SJL-Ptprc$^{a}$ Pepc$^{b}$/BoyJ, cat #002014), TGE600 (cat #020456), and NSG (cat #005557) mice were purchased from Jackson laboratories. Animals were housed in a temperature and humidity-controlled pathogen-free environment and kept under a 12-hour lighting cycle at the Cancer Center Isolation Facility (CCIF), Duke University, NC, USA. Regular welfare checks were made by study investigators and CCIF staff as per IACUC protocols.

#### Murine lymphocyte culturing

Expansion of murine CD3$^{+}$ cells was performed by harvesting spleens from healthy 6–10-week-old genetically identical mice. Spleens were disrupted over a 70-μm strainer before suspension in ACK lysis buffer (Quality Biological, 118-156-101) for 2 minutes. Cells were then re-suspended at a concentration of 2 x 10$^{6}$ cells/mL in complete T cell medium supplemented with either 100 IU/mL IL-2 (ProLeukin), or 20 ng/mL IL-7 (PeproTech, 217-17). Complete T cell medium consists of RPMI 1640 (Sigma-Aldrich, R8758) with 10% FBS and supplemented with 1:100 Minimum Essential Medium Non-Essential Amino Acids (MEM NEAA, Gibco, 1114050), 1:100 Sodium Pyruvate (Gibco, 11360070), 1:100 L-Glutamine (Gibco, 25030081), 1:100 Pen-Strep (Gibco, 15140122), 1:1000 2-Mercaptoethanol (ThermoScientific, 21985023) and 1:1000 Gentamicin (Gibco, 1570060). After collection, cells were stimulated with 2 μg/mL Concanavalin A (Sigma-Aldrich, C5275). Cells were maintained in 24-well tissue culture plates (Corning Falcon) and split every 2-3 days to maintain concentration at 1-2 x 10$^{6}$ cells/mL prior to usage for either ALT or flow cytometry.

#### Leukapheresis and human PBMC culturing

Human PBMCs were obtained by leukapheresis at the Duke Apheresis Unit (SOP-JHS-HDC-CL-023 “Leukapheresis Collection Procedure”) and transported to the Cell Processing facility (SOP-JHS-HDC-CL-011 “Leukapheresis Transport and Receipt Procedure”). Briefly, following leukapheresis, samples were diluted, underlayered with Ficoll (Histopaque-1077, Sigma-Aldrich, 10771), and spun to select for hPBMCs. Samples were then anonymized and stored in liquid nitrogen below -135°C. Cryopreserved PBMCs were then thawed into PrimeXV T-cell expansion media (FujiFilm) supplemented with 3% of human platelet lysate (hPL, Compass Biomedical) and pelleted at 300g for 10 min. The cell pellet was re-suspended in PrimeXV/3% hPL to a concentration of 1 x 10$^{6}$ cells/ml and 1 x 10$^{6}$ cells plated per well of a G-Rex 6M six well plate (Wilson-Wolf). For each sample, two wells were set up. One was supplemented with IL-2 at 300 IU/ml (BioTechne, BT-002-AFL) and the other was supplemented with IL-7 at 20ng/ml (PeproTech, 200-07). The cells were activated with 100µl a-CD3/a-CD8 nanoparticles (TransAct, Miltenyi, 130-111-160) according to the manufacturers protocol. The plates were incubated at 37ºC and 5% CO2. On culture day 3, TransAct was inactivated by adding 10x volume of complete media. On culture day 7, 90% of the media was removed, the cells counted, a small aliquot removed for flow cytometry analysis, the culture split to 1 $x10^6$ cells/$cm^2$ and fresh media added to a total of 100 ml. Cultures were monitored daily for lactate levels and media changed when levels reached 15 mM. On culture day 14, 90% of media was removed, the cells were re-suspended and counted and analyzed. Regarding follow-up experiments evaluating DUSP6 inhibition, naïve hPBMCs were obtained from StemCell (70025.2) and expanded via similar processes. For cells in the DUSP6 inhibition group, culture medium was supplemented with 0.4µM BCI (MedChemExpress, HY-115502, dose based on previously published protocols (1)).

#### Tumor inoculation

All tumor studies in this report placed tumors intracranially in mice and followed protocols described previously (2). Briefly, we collected tumor cells in logarithmic growth phase by first detaching them from culture flasks using 0.05% Trypsin, EDTA (Gibco, 25300-054). Cells were then suspended in sterile phosphate buffered saline (PBS, Gibco, 10010-023) before being mixed in a 1:1 ratio with 10% methylcellulose (Sigma, M-0512). This mixture was loaded into a 250µL Hamilton syringe (Hamilton company, 81120) with a 25-gauge needle attached. Mice were anesthetized with isoflurane before being placed into a stereotactic frame. The scalp was sterilized, and a midline incision made to visualize the bregma. The loaded syringe was positioned 2mm lateral to the right of the bregma and lowered 5mm through the skull, before being retracted 1mm to form a pocket for tumor cell inoculation. 5µL of tumor cells were infused at 120µL per minute. For tumor studies using CT2AvIII, $3 \times 10^4$ cells were infused, studies using U87vIII infused $2.5 \times 10^4$ cells and studies using GL261 infused $6 \times 10^4$ cells per mouse. Tumor infusate was allowed to settle in the pocket for 45 seconds, before withdrawing the needle. The injection site was closed with bone wax and the wound was stapled closed. Mice were allowed to recover before cages were replaced.

#### In vivo adoptive lymphocyte transfer and antibody administration

For adoptive lymphocyte transfer in this study, cells that had been expanded as described in the murine lymphocyte culturing section were collected and suspended in PBS at a concentration of $1 \times 10^8$ cells/mL. 100 µL (equivalent to a dose of $1 \times 10^7$ cells per mouse) was administered via retro-orbital intravenous injection using an Ultra-Fine Lo-Dose insulin syringe (BD, 324703).

Antibody administration of anti-VLA-4 (clone PS/2, BioXCell #BE0071), anti-4-1BB (clone LOB12.3, BioXCell #BP0169), anti-PD-1 (clone 29F.1A12, BioXCell, #BE0273), anti-CTLA-4 (clone 9D9, BioXCell, #BE0164) were performed as part of this study. Intraperitoneal doses consisted of 200 µg antibody made up to 200 µL with PBS and given as described in study descriptions. For intracranial administration (anti-VLA-4), 2 µg antibody was made up to 20 µL with PBS and given in a similar fashion to tumor inoculation above.

hEGFRvIII:CD3 BRiTE (lot #201905) was manufactured at CellDex therapeutics to cGMP standards for a planned clinical trial (NCT04903795) and has been described in detail previously (3). Stock solution of 0.5 mg/mL was maintained at -80°C until use. For administration to animals, 100 µL (equivalent to 50 µg per mouse) was given via retro-orbital intravenous injection using an Ultra-Fine Lo-Dose insulin syringe.

#### Tissue processing and flow cytometry

Procedures for tissue processing and flow cytometry are based on our previously published protocols (4). Briefly, mice were anesthetized with isoflurane before undergoing cardiac puncture and perfusion with PBS. Following perfusion, brain tissue, bone marrow, spleen and lungs were collected and prepared for staining (dependent on experimental design). To isolate tumors, brains were divided into tumor-bearing hemispheres (typically right) and non-tumor-bearing hemispheres (left). Tumor presence was visually confirmed at collection. Blood was collected via retro-orbital eye bleeds or pre-perfusion cardiac puncture. After collection, tissue was homogenized in digestion medium consisting of 0.05mg/mL Liberase DL (Roche, 5401160001), 0.05mg/mL Liberase TL (Roche, 5401020001) and 0.2mg/mL DNAse I (Roche, 10104159001), in HBSS with calcium and magnesium (Gibco, 14175095). Tissue in digestion buffer was placed in a homogenizer (Wheaton, 62400-620) and disaggregated via 5-10 strokes with a loose-fitting (A-size) pestle. This suspension was incubated for 15 minutes at 37°C before filtering through a 70-µm strainer. Note: lung tissue was placed in a shaker for 45 minutes with digestion buffer. Femurs were removed from mice and bone marrow removed by centrifugation at 10,000xg for 30 seconds. After centrifugation, cells were re-suspended in ACK lysis buffer (Quality Biological, 118-156-101) for 2 minutes. Myelin was removed from brain tissue by 30 minutes of centrifugation with 30% Percoll (Sigma Aldrich, P1644) with no brake. After long centrifuge, the myelin layer was carefully aspirated, and the remaining cell pellet resuspended at a concentration of 1-2 x $10^7$ cells/mL in PBS. Analysis of endothelial cells from IC tumor followed published protocols (5). Cells maintained in culture were removed from culture vessels and pelleted at 500g for 5 minutes and similarly resuspended in PBS for flow cytometry.

Before panel staining, Live/Dead Aqua (1:200, ThermoFisher, L34966) was added and co-incubated for 30 minutes at 4°C to assess viability. Brilliant Stain buffer (BD, 563794) was added prior to addition of panels (antibodies listed in Supplementary Table 1). Staining was performed by co-incubation for 30 minutes at 4°C before fixation in flow buffer (PBS, 1% BSA, 0.5 mM EDTA) with 1% formaldehyde. Prior to flow cytometry, counting beads (Invitrogen, C36950) were added to a fixed volume (i.e., 10 μL beads to 300 μL sample in fix buffer). Determination of counts per weight was performed by the following calculations:

$$\text{Cell Concentration (per μL)} = \frac{\text{Number of acquired cells}}{\text{Number of acquired beads}} \text{ x } \frac{\text{Number of beads added to sample}}{\text{Total volume of analyzed sample}}$$

$$\text{Normalized count (per gram)} = \frac{\text{Cell concentration x Total volume of analyzed sample}}{\text{Fraction of sample stained x Tissue weight}}$$

Flow cytometry was performed on a LSR Fortessa (BD) using FACS Diva Software v.9, BD Biosciences and analyzed with FlowJo v.10 (Tree Star). Flow sorting was performed using a Beckman Coulter MoFlo Astrios EQ High Speed Sorter.

#### Cytotoxicity

Tumor cells were labelled with a viability dye (Cell Trace Violet, ThermoFisher, C34557) before being added to a 96-well plate (1 x $10^4$ cells per well). T cells were then added to each well at a 10:1 ratio (1 x $10^5$ cells per well). Antibodies were added at a starting concentration of 10 μg/mL in the first row, before 10-fold serial dilutions were added to subsequent rows. The final row did not have antibody added to act as a control. After 24 hours of co-culture, cells were trypsinized and re-suspended in 115 μL flow buffer with 10 μL of counting beads added per well (Invitrogen, C36950, 125 μL total per well). Plates were analyzed using a high-throughput reader attached to a LSR Fortessa (BD). Percentage viability was assessed based on control well bead-normalized counts representing 100% tumor viability/no antibody effect.

#### Immune functional assays

Immune functional assays were performed with flow-sorted $CD8^+$ human lymphocytes. T cells were either evaluated alone (unstimulated), with tumor (U87vIII) only, with tumor & BRiTE, or with phorbol myristate acetate/ionomycin (1:500). Cells were incubated with GolgiStop (Monensin, BD, 554724) and GolgiPlug (Brefeldin A, BD, 555029) for 6-8 hours at 37C. After incubation, cells were stained for surface markers before being fixed and permeabilized with FixPerm solution (BD,

554714). Cells were then washed in 1x PermWash buffer (BD, 554714) before staining with intracellular antibodies before suspension in 300uL PBS for flow analysis.

#### Immunohistochemistry & toxicity

Toxicity studies were conducted in C57/Bl6 mice. Animals were perfused with PBS as described previously, before brain, spinal cord and liver were removed. Samples were immersed in 10% neutral buffered formalin (NBF) for 72 hours before being moved to 70% ethanol. Following fixation, samples were formalin-fixed paraffin-embedded at HistoWiz inc., before being sectioned (5-micron) and stained with Hematoxylin & Eosin (H&E) and Luxol Fast Blue (LFB). Slides were reviewed by veterinary pathologists. Serum was collected at termination for clinical chemistry and analyzed on a Heska veterinary chemistry analyzer within the Duke Division of Laboratory Animal Resources (DLAR).

#### RNA sequencing assays and analysis

Flow-sorted $CD8^+$ T cells were snap frozen in cell pellets and analyzed via bulkRNA sequencing at Azenta life sciences. Principal Component Analysis (PCA) performed after normalization of raw gene counts using the logCPM method, before filtering the most variable 2500 genes (transformed using the Z-score method). PCA analysis was generated using BioJupies (6). Hierarchical cluster heatmaps were generated via the same method and using clustergrammer (Maayan lab, (7)). Gene Ontology (GO) biological processes analysis was performed using the NIH database for annotation, visualization and integrated discovery dashboard (DAVID (8)). Volcano plots were generated based on tests for differential expression (Wald test used to generate p-values and log2 fold changes) created with DESeq2 (9) and visualized using VolcaNoseR (10).

### Supplemental Figures

#### Supplemental Figure 1

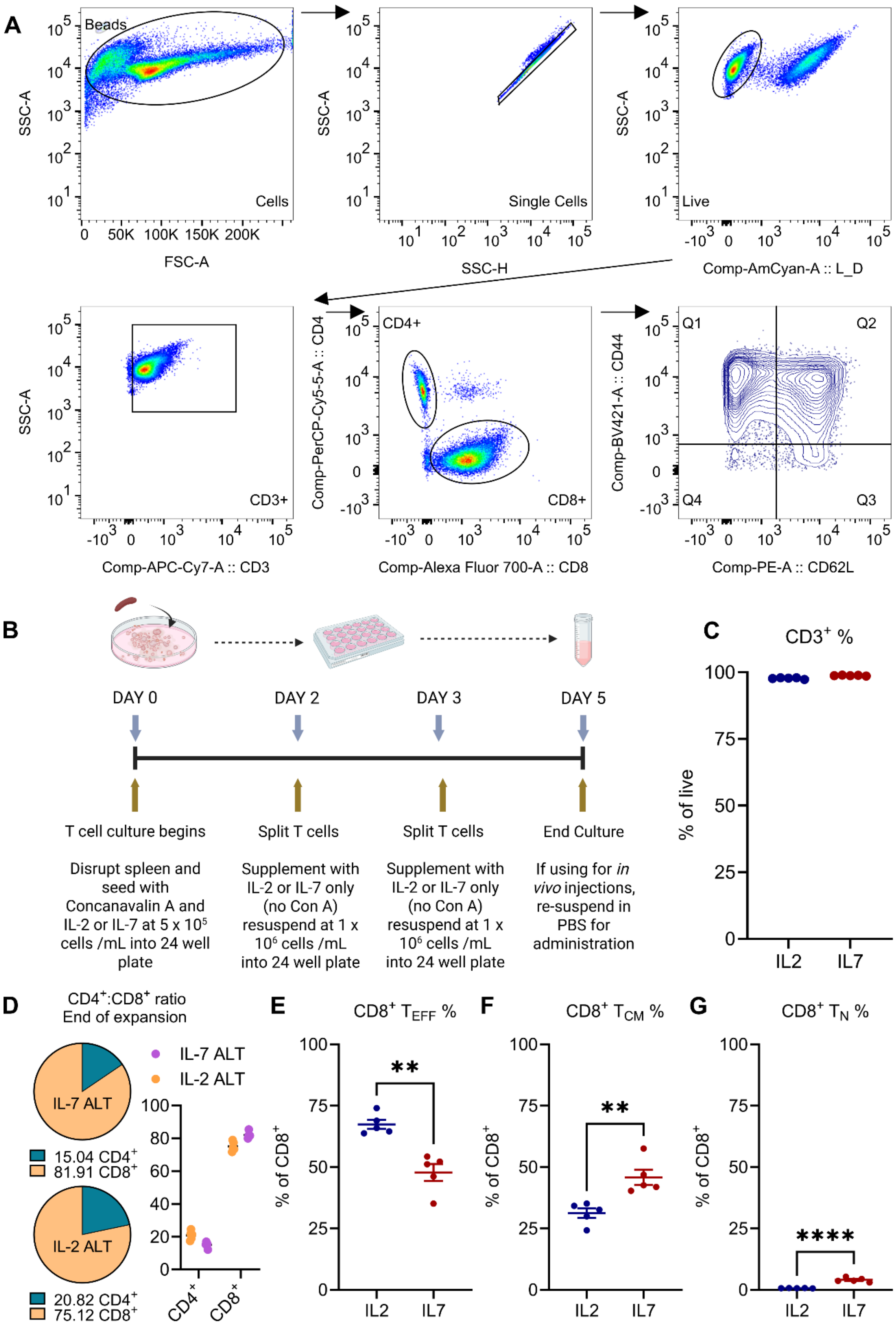

**(A)** Representative gating strategy shown for phenotypic analysis of cellular product at expansion end.

**(B)** Overview of *ex vivo* mouse lymphocyte culturing process.

**(C)** $CD3^+$ fraction at expansion end for both IL-2 and IL-7 co-culture conditions (n=5/group for all graphs).

**(D)** Pie-charts showing CD4: CD8 fraction at expansion end, with dot-plot showing individual values across groups.

**(E-G)** $CD8^+$ $T_{EFF}$, $CD8^+$ $T_{CM}$ and $CD8^+$ $T^N$ fractions at expansion end.

Statistical analyses performed using unpaired T tests and data presented as mean ± SEM unless otherwise specified. Experimental outlines generated using BioRender.com.

Supplemental Figure 2

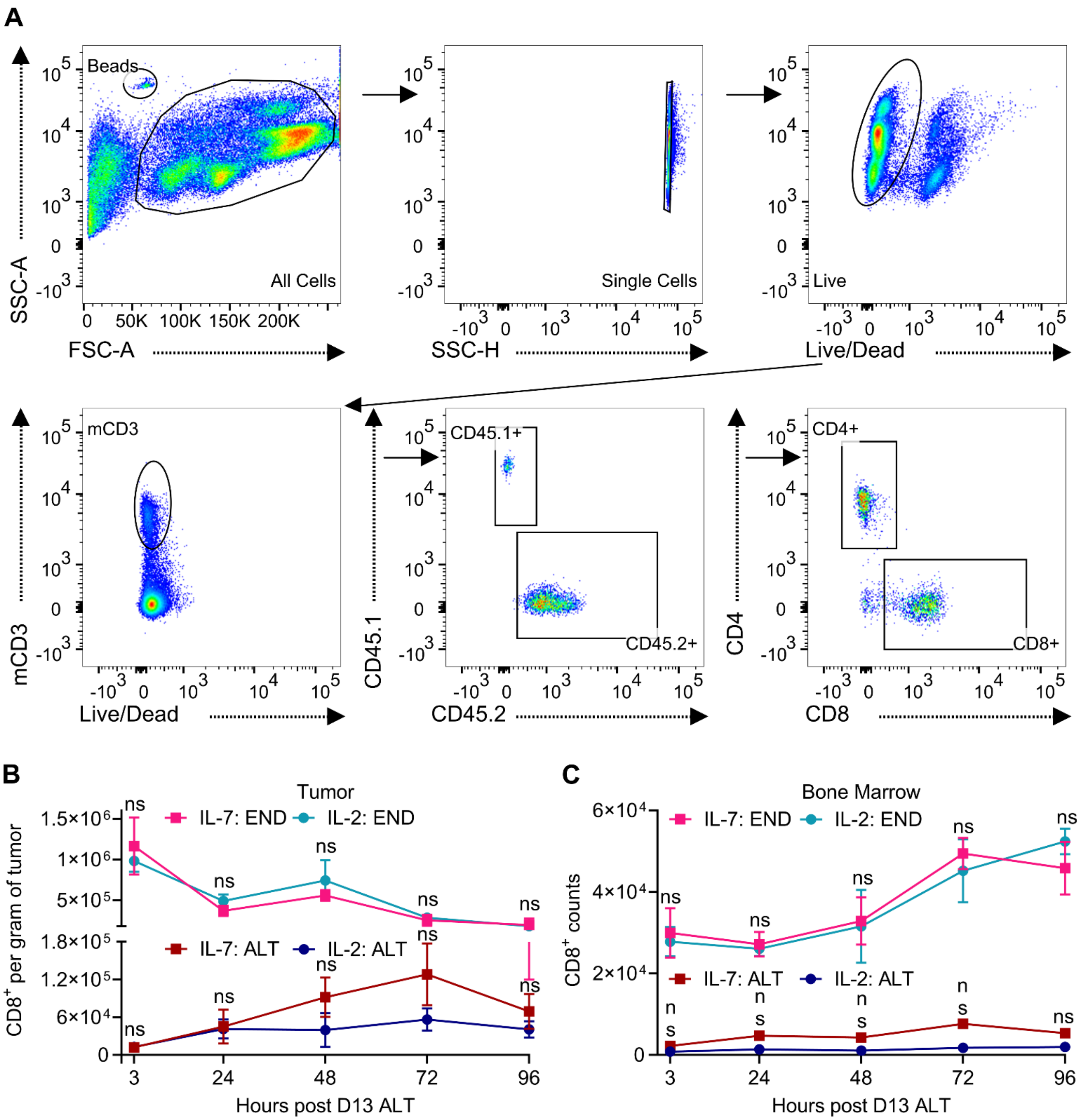

**(A)** Representative gating strategy shown for analysis of CD45.1$^+$CD8$^+$ ALT during time course experiment.

**(B)** Weight-normalized counts of exogenous CD45.1$^+$CD8$^+$ entry into tumor over time (n=4-5/group for all graphs).

**(C)** Counts for endogenous and exogenous CD45.1$^+$CD8$^+$ entry into bone marrow over time.

Statistical analyses performed via two-way ANOVA and data presented as mean ± SEM unless otherwise specified.

#### Supplemental Figure 3

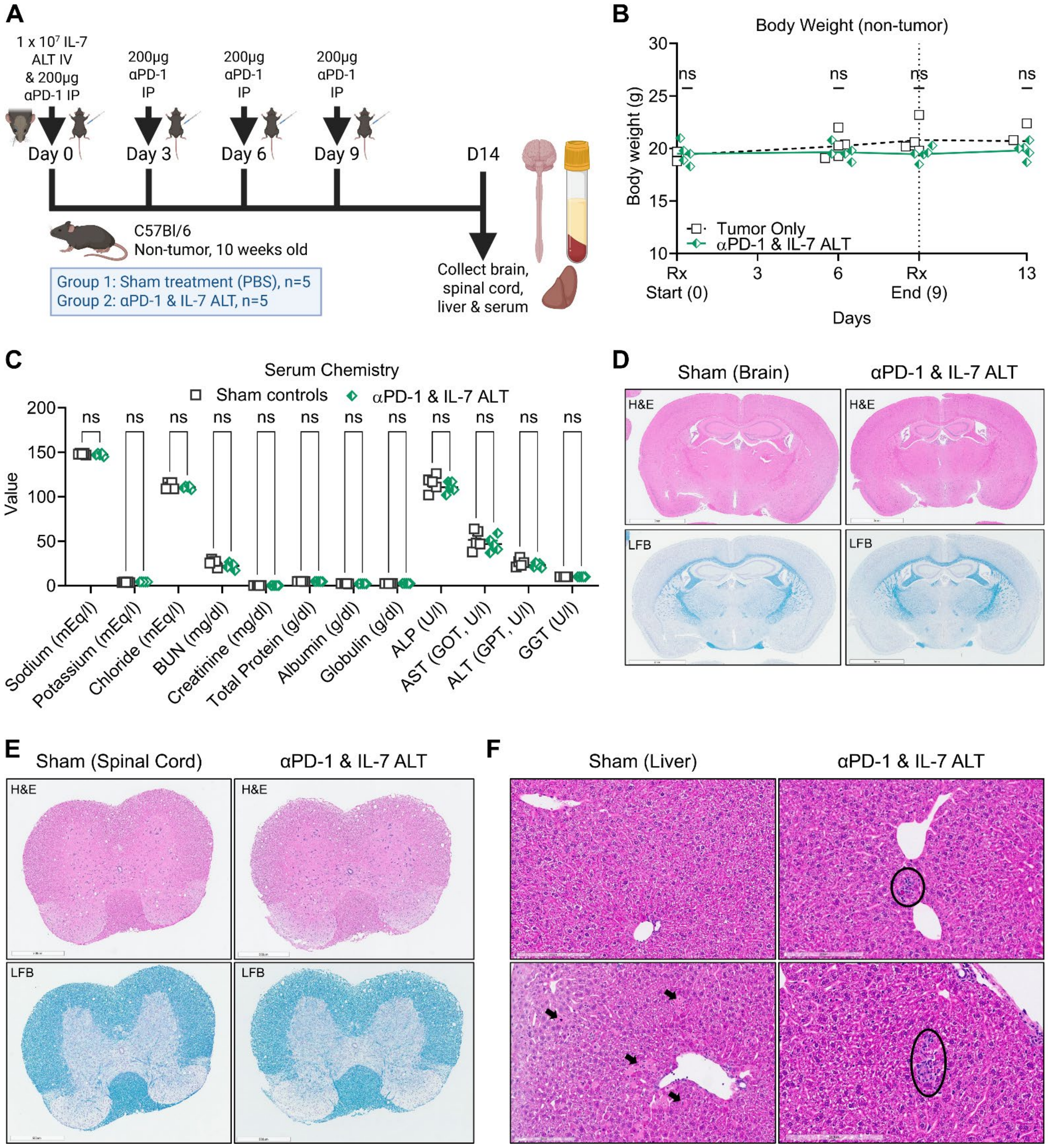

**(A)** Study overview of toxicity study in non-tumor female C57BL/6 mice (n=5/group for all graphs).

**(B)** Bodyweight monitoring for mice throughout the treatment course and for 5 days following treatment end.

**(C)** Serum chemistry values including electrolytes and liver function tests for mice treated with combination αPD-1 & IL-7 ALT vs sham (D14). No significant differences were observed.

**(D & E)** Representative photomicrographs of brain (scalebar: 3mm) and spinal cord (scalebar: 500μm) from both groups. H&E staining shown on top rows, Luxol Fast Blue on bottom rows.

**(F)** Representative H&E photomicrographs of liver (scalebar: 200μm). Sham column shows healthy liver (top left) along with hepatocellular degeneration and necrosis without associated inflammation (post-mortem changes, arrows, bottom left). Combination αPD-1 & IL-7 ALT therapy column with small foci of hepatocellular necrosis with associated mixed infiltrates (circled, top and bottom right).

Statistical analyses performed using multiple unpaired t-tests and data presented as mean ± SEM.

Supplemental Figure 4

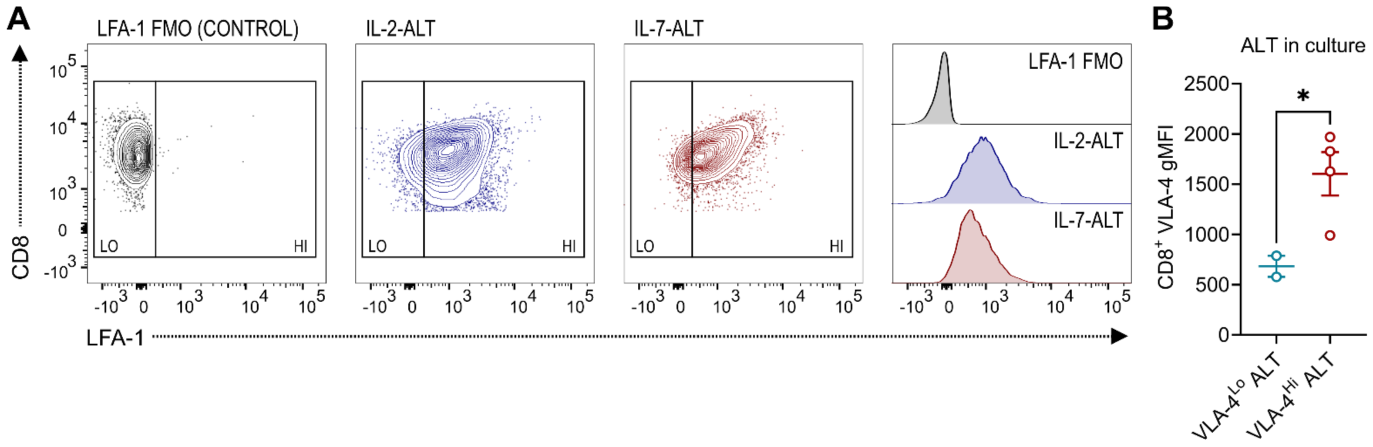


**(A)** Gating strategy for CD8⁺ LFA-1.

**(B)** Pooled analyses of lymphocyte expansions (representative of 3 experiments) with IL-7 or IL-2 to produce ALT which was VLA-$4^{Hi}$ VLA-$4^{Lo}$ (VLA-$4^{Hi}$ n=4 vs VLA-$4^{Lo}$ n=2, unpaired t-test).

#### Supplemental Figure 5

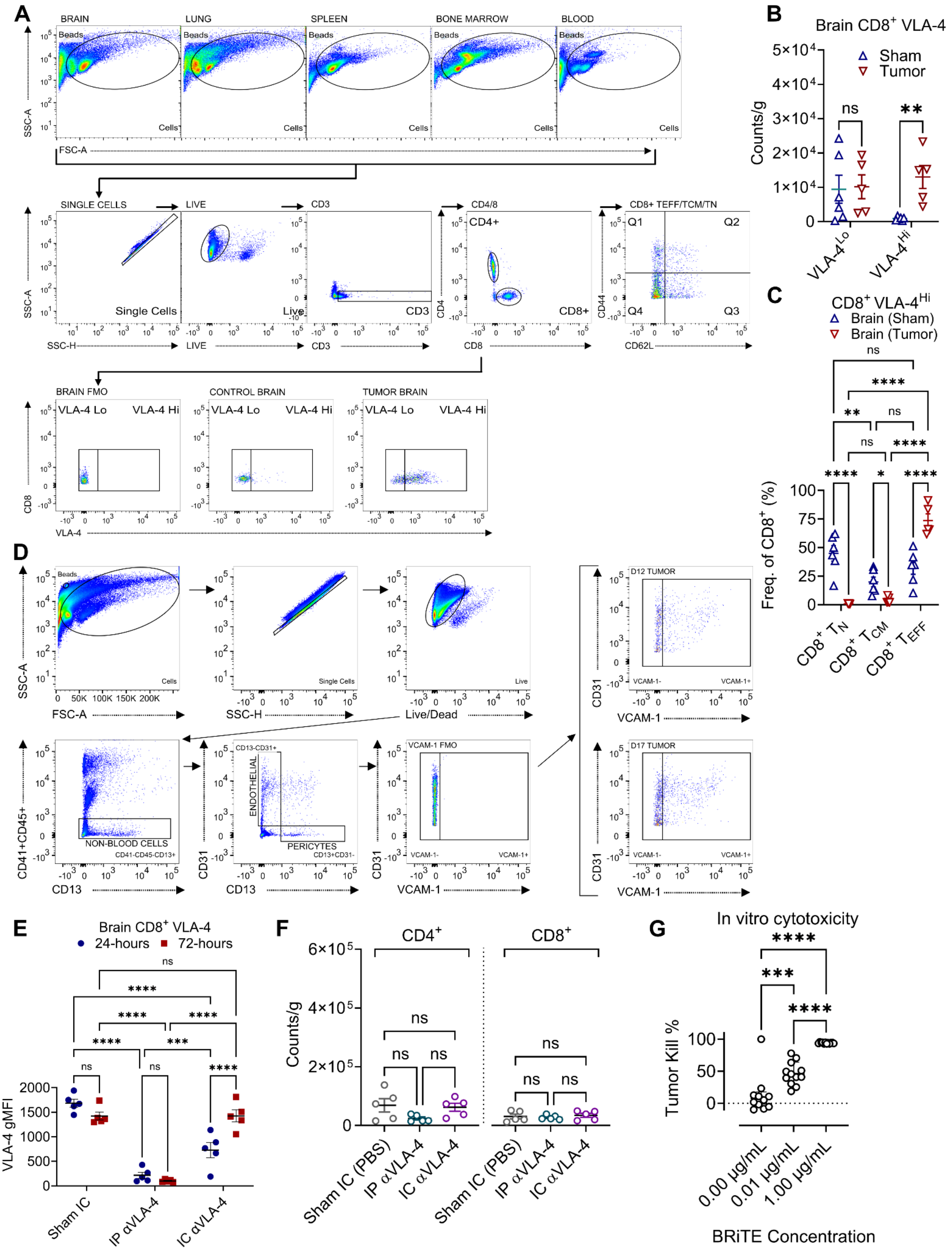

**(A)** Representative gating strategy for VLA-4 compartment experiment.

**(B)** $CD8^+$ T cell weight adjusted counts for the $VLA\text{-}4^{Lo}$ and $VLA\text{-}4^{Hi}$ populations (n=5-6/group)**.** Comparisons via multiple unpaired t-tests.

**(C)** Phenotypic make up of $CD8^+VLA\text{-}4^{Hi}$ cells in sham controls/tumors.

**(D)** Representative gating strategy for CNS endothelial/pericyte VCAM-1.

**(E)** $CD8^+$ VLA-4 gMFI on T cells in tumors 24- and 72-hours following sham injections, intraperitoneal αVLA-4 (200μg) or intracranial αVLA-4 (2μg) administration.

**(F)** IC $CD4^+$ & $CD8^+$ counts 24-hours following administration of IP/IC αVLA-4 or sham controls. Comparisons via one-way ANOVA.

**(G)** In vitro cytotoxicity assay with BRiTE to establish half-maximal effective concentration, $EC_{50}$ of 0.01μg/mL (n=12/group). Comparisons shown via one-way ANOVA.

Statistical analyses performed using two-way ANOVA and data presented as mean ± SEM unless otherwise specified.

#### Supplemental Figure 6

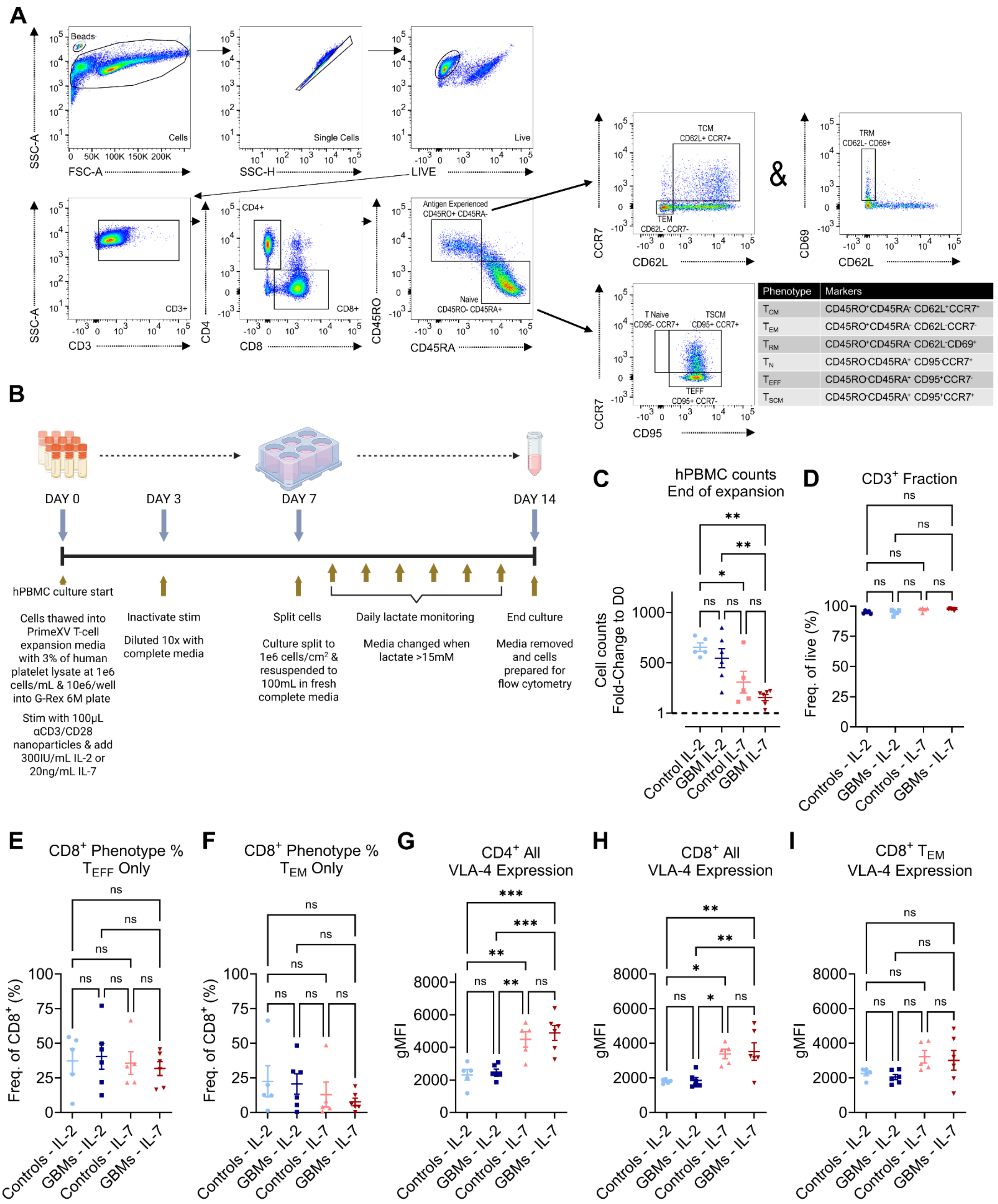


| Phenotype | Markers |
|---|---|
| $T_{CM}$ | $CD45RO^+CD45RA^- CD62L^+CCR7^+$ |
| $T_{EM}$ | $CD45RO^+CD45RA^- CD62L^-CCR7^-$ |
| $T_{RM}$ | $CD45RO^+CD45RA^- CD62L^-CD69^+$ |
| $T_N$ | $CD45RO^-CD45RA^+ CD95^-CCR7^+$ |
| $T_{EFF}$ | $CD45RO^-CD45RA^+ CD95^+CCR7^-$ |
| $T_{SCM}$ | $CD45RO^-CD45RA^+ CD95^+CCR7^+$ |

**(A)** Representative gating strategy shown for phenotyping of cellular product.

**(B)** Overview of *ex vivo* hPBMC culturing process.

**(C)** hPBMC fold-increase comparisons across all 4 groups at expansion end (n=5-6/group for all graphs).

**(D)** $CD3^+$ fraction at expansion end.

**(E & F)** $CD8^+$ $T_{EFF}$ and $CD8^+$ $T_{EM}$ fractions at expansion end.

**(G)** $CD4^+$ VLA-4 geometric mean fluorescence intensity shown across all 4 conditions.

**(H)** $CD8^+$ VLA-4 geometric mean fluorescence intensity across all 4 conditions.

**(I)** $CD8^+$ $T_{EM}$ VLA-4 geometric mean fluorescence intensity across all 4 conditions.

Comparisons via one-way ANOVA and data presented as mean ± SEM unless otherwise specified. Experimental outlines generated using BioRender.com.

#### Supplemental Figure 7

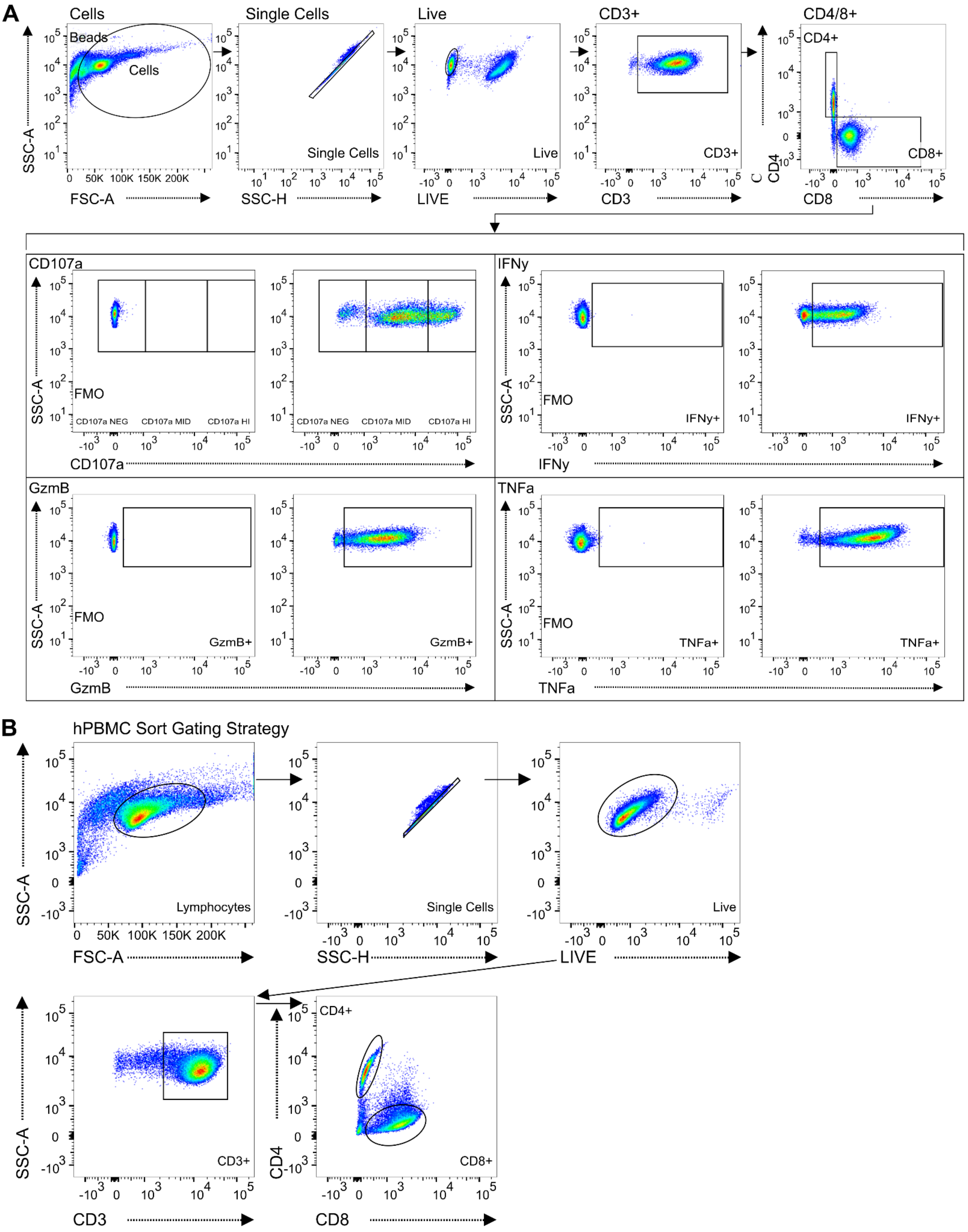

**(A)** Flow gating strategy shown for $CD107a^{Hi}$, interferon-gamma (IFN-*γ*), Granzyme B (GzmB) and tumor-necrosis factor-alpha (TNF-*α*).

**(B)** Flow gating strategy for flow-sorting of $CD8^+$ T cells from end expansion product of hPBMCs from patients with glioblastoma.

#### Supplemental Figure 8

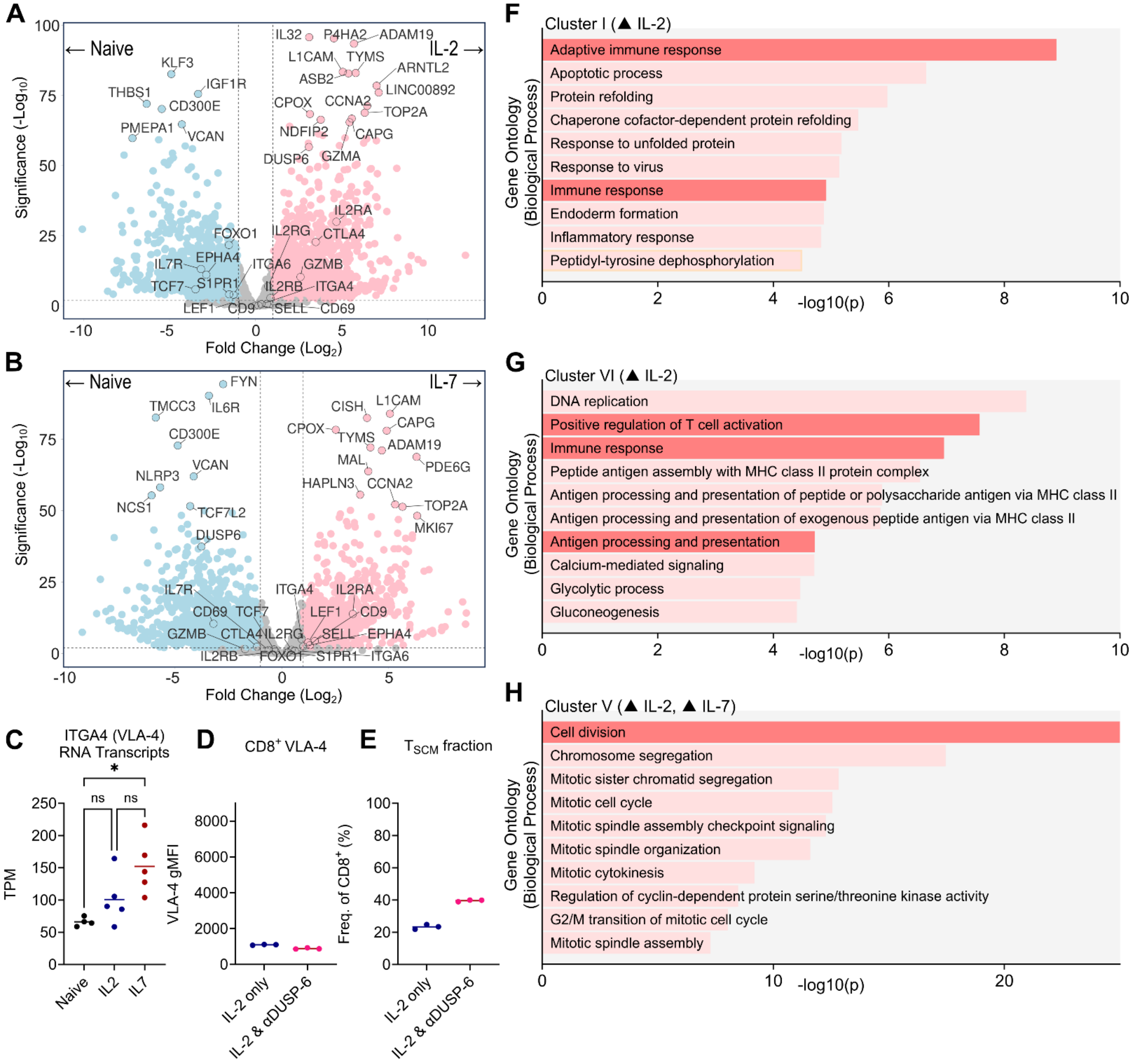

**(A & B)** Volcano plots comparing differential gene expression between naïve and IL-2 or naïve and IL-7 expanded $CD8^+$ T cells.

**(C)** Comparisons of raw transcripts per million (TPM) across naïve, IL-2-expanded or IL-7-expanded groups for *ITGA4* (VLA-4).

**(D)** $CD8^+$ VLA-4 expression (gMFI) when culturing with IL-2 alone, or IL-2 supplemented with 0.4μm αDUSP-6 inhibitor (BCI).

**(E)** $CD8^+$ $T_{SCM}$ fraction when culturing with IL-2 alone, or IL-2 supplemented with 0.4μm αDUSP-6 inhibitor (BCI).

**(F & G)** Gene Ontology biological processes analysis of genes expressed in cluster I and VI (IL-2 upregulated, *GZMA, GZMB, IFNγ, FASLG, CD69, CTLA-4, TOX* etc.)

**(H)** Gene Ontology biological processes analysis of genes expressed in cluster V (upregulated in IL-2 and IL-7, *MKI67* etc.)

Volcano plots were generated using VolcaNoseR (10). Gene Ontology (GO) biological processes analysis was performed using the NIH DAVID database (8). Comparisons via one-way ANOVA and data presented as mean ± SEM unless otherwise specified.

### Supplemental Tables

#### Supplemental Table 1. Overview of mouse flow cytometry antibodies

| PHENOTYPING-1 | | | | | | |
|---|---|---|---|---|---|---|
| **Antigen** | **Company** | **Clone** | **Fluorophore** | **Catalog #** | **Vol/1e6 cells (μL)** | **Purpose** |
| CD45.1 | Biolegend | A20 | APC | 110714 | 1 | T Cell trafficking |
| CD4 | Biolegend | RM4-5 | PerCp-Cy5.5 | 100540 | 0.5 | CD4+ (Helper) T |
| CD8a | BD | 53-6.7 | APC-H7 | 560182 | 3 | CD8+ (Cytotoxic) T |
| CD44 | BD | IM7 | BV786 | 563736 | 1 | Memory |
| CD62L | BD | MEL-14 | BB515 | 565261 | 0.5 | Memory |
| VLA-4 | Biolegend | R1-2 | PE-Dazzle 594 | 103626 | 1 | Integrin |
| LFA-1 | BD | M18/2 | BV421 | 744597 | 3 | Integrin |
| CD45.2 | BD | 104 | BV605 | 563051 | 3 | T cell trafficking |
| PD-1 | Biolegend | 29F.1A12 | PE | 135205 | 1 | Exhaustion |
| CD3 epsilon | Biolegend | 145-2C11 | BV711 | 100349 | 1 | T -Cells |
| **PHENOTYPING-2** | | | | | | |
| **Antigen** | **Company** | **Clone** | **Fluorophore** | **Catalog #** | **Vol/1e6 cells (μL)** | **Purpose** |
| CD3 | Biolegend | 17A2 | APC-Cy7 | 100222 | 1 | T-Cells |
| CD4 | Biolegend | GK1.5 | PerCP-Cy5.5 | 100434 | 3 | CD4+ (Helper) T |
| CD8 | Biolegend | 53-6.7 | AF700 | 100730 | 5 | CD8+ (Cytotoxic) T |
| CD45.1 | Biolegend | A20 | BV785 | 110743 | 3 | Exogenous |
| CD45.2 | Biolegend | 104 | PE-Cy7 | 109830 | 1 | Endogenous |
| CD44 | Biolegend | IM7 | BV421 | 103040 | 1 | Memory |
| CD62L | Biolegend | MEL-14 | PE | 104408 | 3 | Memory |
| CD69 | Biolegend | H1.2F3 | AF647 | 104518 | 3 | Activation |
| CD25 | Biolegend | PC61 | BV711 | 102049 | 3 | Activation |
| CD49d (VLA-4) | Biolegend | R1-2 | PE-Dazzle 594 | 103626 | 3 | Integrin |
| CD18 (LFA-1) | BD | C71/16 | BV650 | 740465 | 1 | Integrin |

| ENDOTHELIAL (VCAM-1) EXPRESSION | | | | | | |
|---|---|---|---|---|---|---|
| **Antigen** | **Company** | **Clone** | **Fluorophore** | **Catalog #** | **Vol/1e6 cells (µL)** | **Purpose** |
| CD31 | BD | MEC 13.3 | APC | 551262 | 20 | Endothelial cells |
| CD13 | BD | R3-242 | FITC | 558744 | 8 | Pericytes |
| CD41 | BD | MWReg30 | PE | 558040 | 4 | Negative Selection |
| CD45 | BD | 30-F11 | PE | 553081 | 4 | Negative Selection |
| VCAM-1 | BD | 429 MVCAM.A | BV786 | 740865 | 8 | Endothelial binding |

Viability assessed using LIVE/DEAD™ Fixable Aqua Dead Cell Stain Kit, ThermoFisher, Catalog Number #L34966

#### Supplemental Table 2. Overview of human flow cytometry antibodies

| PHENOTYPING | | | | | | |
|---|---|---|---|---|---|---|
| **Antigen** | **Company** | **Clone** | **Fluorophore** | **Catalog #** | **Vol/1e6 cells (μL)** | **Purpose** |
| CD3 | BD | SK7 | APC-H7 | 560176 | 3 | T-Cells |
| CD4 | Biolegend | SK3 | PerCP-Cy5.5 | 344608 | 1 | CD4+ (Helper) T |
| CD8 | Biolegend | SK1 | AF700 | 344724 | 1 | CD8+ (Cytotoxic) T |
| CD45RA | BD | HI100 | BB515 | 564552 | 0.5 | Memory |
| CD45RO | Biolegend | UCHL1 | BV605 | 304238 | 3 | Memory |
| CD62L | Biolegend | DREG-56 | PE | 304806 | 1 | Memory |
| CCR7 | Biolegend | G043H7 | BV421 | 353208 | 5 | Memory |
| CD95 | Biolegend | DX2 | PE-Cy7 | 305622 | 1 | Fas |
| CD69 | Biolegend | FN50 | BV785 | 310932 | 3 | Early Activation |
| HLA-DR | Biolegend | L243 | APC | 307610 | 5 | Activation |
| CD49d (VLA-4) | Biolegend | 9F10 | PE-Dazzle 594 | 304326 | 1 | Integrin |
| CD18 (LFA-1) | BD | L130 | BV650 | 744553 | 5 | Integrin |
| **FLOW SORT AND IMMUNE FUNCTIONAL ASSAYS** | | | | | | |
| **Antigen** | **Company** | **Clone** | **Fluorophore** | **Catalog #** | **Vol/1e6 cells (μL)** | **Purpose** |
| CD3 | Biolegend | SK7 | PE-Cy7 | 344816 | 5 | T-Cells |
| CD4 | Biolegend | SK3 | PerCP-Cy5.5 | 344608 | 5 | CD4+ (Helper) T |
| CD8 | Biolegend | SK1 | AF700 | 344724 | 5 | CD8+ (Cytotoxic) T |
| CD107a | Biolegend | H4A3 | PE | 328607 | 5 | Degranulation |
| IFN-y | Biolegend | 4S.B3 | BV650 | 502537 | 5 | Function |
| GzmB | Biolegend | QA16A02 | APC | 372203 | 5 | Cytotoxic Function |
| TNF-a | Biolegend | MAb11 | BV605 | 502935 | 5 | Function |

Viability assessed using LIVE/DEAD™ Fixable Aqua Dead Cell Stain Kit, ThermoFisher, Catalog Number #L34966
